# A condensate forming tether for lariat debranching enzyme is defective in non-photosensitive trichothiodystrophy

**DOI:** 10.1101/2022.11.03.515106

**Authors:** Brittany A. Townley, Luke Buerer, Albino Bacolla, Timur Rusanov, Nicolas Schmidt, Sridhar N. Srivatsan, Nathanial E. Clark, Fadhel Mansoori, Reilly A. Sample, Joshua R. Brickner, Drew McDonald, Miaw-Sheue Tsai, Matthew J. Walter, David F. Wozniak, Alex S. Holehouse, John A. Tainer, William G. Fairbrother, Nima Mosammaparast

## Abstract

The pre-mRNA life cycle requires intron processing; yet, how intron processing defects influence splicing and gene expression is unclear. Here, we find TTDN1, which is frequently mutated in non-photosensitive trichothiodystrophy (NP-TTD), functionally links intron lariat processing to the spliceosome. The conserved TTDN1 C-terminal region directly binds lariat debranching enzyme DBR1, while its N-terminal intrinsically disordered region (IDR) binds the intron binding complex (IBC). The IDR forms condensates *in vitro* and is needed for IBC interaction. TTDN1 loss causes significant intron lariat accumulation, as well as splicing and gene expression defects, mirroring phenotypes observed in NP-TTD patient cells. *Ttdn1^Δ/Δ^* mice recapitulate intron processing defects and neurodevelopmental phenotypes seen in NP-TTD. A DBR1-IDR fusion recruits DBR1 to the IBC and circumvents the requirement for TTDN1, indicating this tethering role as its major molecular function. Collectively, our findings unveil key functional connections between lariat processing, splicing outcomes, and NP-TTD molecular pathology.

## Introduction

Eukaryotic gene expression involves the recruitment of transcriptional machinery to the transcriptional start site followed by the release of RNA Polymerase II (Pol II), leading to Pol II elongation and the subsequent synthesis of pre-mRNA. While the removal of introns from a nascent pre-mRNA molecule can occur as a post-transcriptional regulatory step, co-transcriptional splicing is an essential feature of many highly-expressed genes (Merkhofer et al., 2014; Naftelberg et al., 2015; Wissink et al., 2019). The ability of cells to undergo co-transcriptional splicing centers on the idea that spatiotemporal organization of splicing and transcription not only protects nascent RNAs from degradation, but also enhances local substrate concentration, thereby increasing reaction efficiencies (Neugebauer, 2019). This subnuclear coordination is facilitated by interactions between the RNA Pol II carboxy-terminal domain (CTD) and a host of factors that regulate transcription, pre-mRNA splicing, mRNA capping, polyadenylation, and downstream steps such as mRNA export (Custodio and Carmo-Fonseca, 2016; Harlen and Churchman, 2017; Hirose and Manley, 1998; Hirose et al., 1999; Ho and Shuman, 1999; Zaborowska et al., 2016). Aside from physically promoting splicing, changes in Pol II elongation rate can broadly influence alternative splicing patterns. Studies examining elongation rate in response to UV damage, chromatin state, and Pol II mutation have found that abnormal elongation rates caused by altered Pol II function reduced splicing efficiency and resulted in aberrant alternative splicing patterns (Dujardin et al., 2014; Fong et al., 2014; Hnilicova et al., 2013; Luco et al., 2011; Munoz et al., 2009; Schor et al., 2013).

While most research on Pol II elongation has centered on the downstream effects on splicing, multiple studies provide evidence for a positive feedback mechanism between early-stage spliceosome assembly and efficient Pol II elongation. Rapid inactivation of the U2 snRNP, an essential spliceosomal component, via small-molecule inhibition was found to largely prevent the release of paused Pol II into the gene body for active transcription elongation, resulting in a global decrease in mRNA biogenesis (Caizzi et al., 2021). In *S. cerevisiae*, blocking pre-spliceosome complex formation via depletion of the RNA helicase Prp5p leads to Pol II accumulation on introns and decreased elongation within intron-containing genes, while transcription of intronless genes is unaffected (Chathoth et al., 2014). In a similar fashion, depletion of the serine/arginine-rich (SR) protein SC-35 in mouse embryonic fibroblasts results in gene-specific Pol II elongation defects (Lin et al., 2008). That RNA Pol II physically interacts with early components of the spliceosome (Zhang et al., 2021) further suggests an intricate interplay between nascent RNA production and its downstream processing.

Although early inhibition of splicing has a demonstrated influence on nascent transcription, whether late-stage splicing inhibition may result in similar alterations is unknown. Notably, a largely understudied terminal step in splicing occurs upon exon ligation, as introns that are removed from the pre-mRNA transcript form a circular RNA fragment known as a lariat. The intron lariat circularizes via a 2’,5’-phosphodiester bond, and the lariat is subsequently linearized by the highly conserved RNA debranching metalloenzyme (DBR1) (Cheng and Menees, 2011; Clark et al., 2016; Montemayor et al., 2014). Human genes have an average of 7-8 introns, and spliceosome assembly occurs de novo on each intron of a pre-mRNA transcript, necessitating an efficient and accurate method of intron removal (Sakharkar et al., 2005; Wahl et al., 2009). Known as the intron lariat turnover pathway, this late-stage step is critical for the release and processing of a subset of regulatory microRNAs (Okamura et al., 2007; Ouchane et al., 1995), as well as for the recycling of spliceosome-associated small nuclear ribonucleoproteins (snRNPs) (Hirose et al., 2003). In the absence of efficient lariat processing, retention of snRNPs in lariat-containing, late-stage splicing complexes has been reported to reduce the efficiency of subsequent spliceosome assembly (Han et al., 2017). Differential regulation of spliceosome-associated snRNP levels influences alternative splicing during normal development and across cancer subtypes (Dvinge et al., 2019). Yet how the rate at which released and recycled intron-associated splicing factors and snRNPs influence gene expression is largely unstudied. Importantly, the consequences of disrupted lariat complex processing, including pleiotropic developmental defects and increased susceptibility to viral infection, imply that this end-stage splicing step has key homeostatic roles in the cell (Li et al., 2016; Zhang et al., 2018). However, it has yet to be determined whether these are a direct effect of increased RNA lariats or an indirect impact on transcription or RNA processing.

Here, we identify the uncharacterized protein TTDN1 as an unappreciated physical and functional link between intron metabolism and gene expression. We show that TTDN1 tethers DBR1 to active splicing complexes to promote the processing of nascent intron lariats, and that lariat accumulation in the absence of TTDN1 coincides with length-dependent gene expression changes. Mutations to the gene encoding TTDN1 constitute a majority of non-photosensitive trichothiodystrophy (NP-TTD) cases, which feature broad neurological and developmental abnormalities thought to be associated with transcriptional defects (Botta et al., 2007; Faghri et al., 2008; Kuschal et al., 2016). We validate our *in vitro* findings by developing a *Ttdn1^Δ/Δ^* mouse model, which recapitulates the RNA processing defects seen in NP-TTD patient cell lines, as well as the neurodevelopmental phenotypes seen in NP-TTD patients. Collectively, our work connects disrupted lariat processing to downstream consequences on gene expression outcomes in the context of the molecular pathology underlying NP-TTD.

## Results

### TTDN1 contains a highly disordered prion-like domain

We recently characterized a link between alkylation damage responses and *RNF113A*, a gene associated with NP-TTD (Brickner et al., 2017; Corbett et al., 2015; Mendelsohn et al., 2020; Tessarech et al., 2020; Tsao et al., 2021). We were curious about the molecular mechanism of pathology associated with *TTDN1*, which is the most commonly altered gene associated with repair-proficient TTD (Faghri *et al*., 2008). While little is known about the structure of TTDN1, previous work defined two C-terminal regions of TTDN1 conserved in metazoans, conserved regions 1 and 2 (CR1 and CR2) (Nakabayashi et al., 2005), the latter of which is the site of a common mutation, M144V, associated with trichothiodystrophy (**Figure 1a**, asterisk). Upon closer inspection of the TTDN1 sequence, we noticed the N-terminal region consists of a 122-residue aromatic-rich prion-like domain with evenly spaced aromatic residues (**Figure 1a**, aromatics in orange). Aromatic-rich prion-like domains have been implicated as drivers of biomolecular condensates *in vitro* and *in vivo* (Kato et al., 2012; Wang et al., 2018).

**Figure 1.**
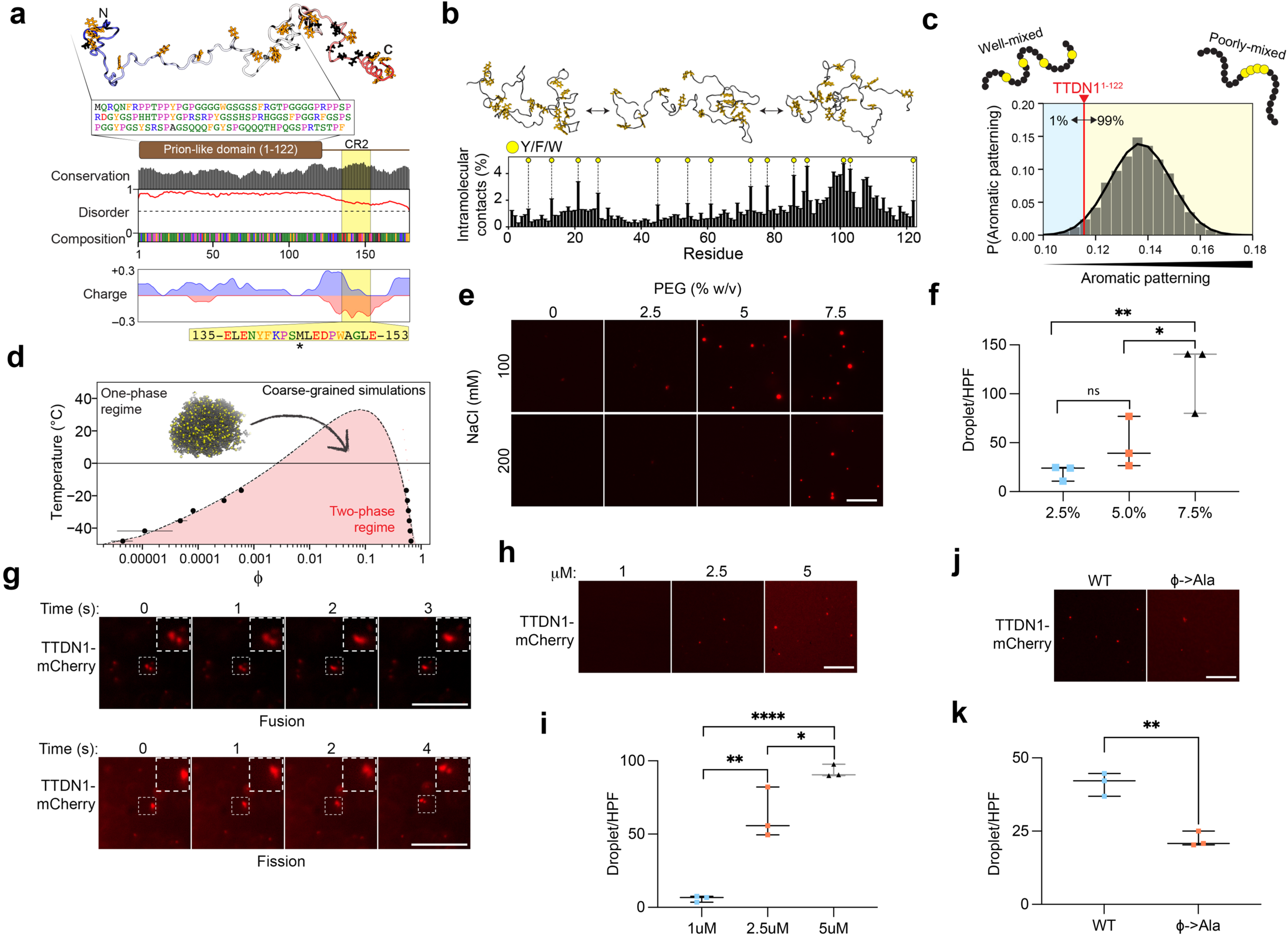
TTDN1 N-terminal domain contributes to condensate formation *in vitro.* **(a)** Schematic of human TTDN1 highlighting its prion-like domain and its conserved CR2 region. **(b)** Contact order analysis was used to quantify the percentage of the simulation in which a given residue is in direct intramolecular contact with any other residue. **(c)** Aromatic patterning was quantified as described previously, generating 10^6^ random sequence permutations to build the null-model distribution (grey histogram) (Martin *et al*., 2020). **(d)** Coarse-grained simulations were run as a function of temperature and the resulting phase boundaries show good agreement with a hypothetic phase diagram generated using Flory-Huggins theory. Temperature is converted from simulation temperature to Celsius using a previously defined scaling factor. Error bars are the standard error of the mean as calculated over three independent simulations. **(e)** Fluorescence microscopy images of TTDN1-mCherry droplets (2.5 µM) under the indicated conditions. Scale bar, 10 μm. **(f)** Droplet quantification per high powered field (HPF) from **(e)**. N = 3 independent replicates per condition. **(g)** Examples of fusion (top) and fission (bottom) of TTDN1-mCherry condensates. Scale bar, 20 μm. **(h)** Microscopy of TTDN1-mCherry droplets at the indicated protein concentrations (200 mM NaCl, no PEG). Scale bar, 10 μm. **(i)** Droplet quantification of **(h)** as in **(f)**. **(j)** Fluorescence microscopy images of TTDN1-mCherry and TTDN1^φ→Ala^-mCherry (2.5 μM) under identical conditions (300 mM NaCl, 5% PEG). Scale bar, 10 μm. **(k)** Droplet quantification of **(j)** as in **(f)**. N = 3 independent replicates per condition. Scale bar, 10 μm. For all droplet quantifications, * *p* < 0.05, ** *p* < 0.01, **** *p* < 0.0001, n.s., not significant, by unpaired t-test.

As such, we sought to develop molecular insight into how TTDN1 might behave in solution. All-atom Monte Carlo simulations of this N-terminal disordered region (TTDN1^1-122^) revealed a heterogeneous and broad distribution of conformations consistent with a disordered protein that engages at most in transient intramolecular interactions (**Supplemental Figure S1a**). With the possible exception of low levels of transient helicity in residues 90-110, secondary structure analysis confirmed the disordered nature of the N-terminus (**Supplemental Figure S1b and Supplemental Video S1**). Given the similarity of TTDN1^1-122^ to other disordered regions studied using similar approaches (Martin et al., 2020), the simulation results support a model in which TTDN1^1-122^ is entirely unstructured. Despite the absence of persistent residual structure, we wondered if distinct residues played an outsized role in the conformational ensemble of TTDN1^1-122^. We performed contact analysis to quantify the extent to which each residue engages in intra-molecular interactions, and identified aromatic residues as the mediators of transient but well-defined intramolecular contacts (**Figure 1b**, yellow circles). We calculated that the natural patterning of these aromatic residues in TTDN1^1-122^ is more evenly distributed than almost all possible patterns of aromatic residues obtained by random chance, implying strong evolutionary pressure for their uniform spacing, in line with prior work on prion-like domains that undergo phase separation (Martin *et al*., 2020) (**Figure 1c**).

Well-distributed aromatic residues that engage in transient intramolecular interactions have been shown to undergo phase separation *in vitro* and *in vivo* (Martin *et al*., 2020; Wang *et al*., 2018). To explore the idea that the N-terminal aromatics residues may function in liquid-liquid phase separation (LLPS) by TTDN1, we applied a previously-developed simple stickers-and-spacers coarse-grained model to quantify the phase behavior of TTDN1^1-122^ (Martin *et al*., 2020). These simulations predicted that TTDN1^1-122^ can undergo phase separation in a temperature-dependent manner (**Figure 1d**). Intriguingly, the driving force for phase separation in TTDN1^1-122^ is predicted to be weaker than that of low-complexity disordered regions studied using the same approach (**Supplemental Figure S1c,** (Martin *et al*., 2020)). While TTDN1^1-122^ shows many of the molecular features one might associate with a protein that robustly drives phase separation (a ‘scaffold’ protein), our simulation data suggest it may behave more as a ‘client’, partitioning into pre-existing biomolecular condensates, as opposed to driving the formation of *de novo* condensates through homotypic interactions. In agreement with this, intramolecular scaling map simulations revealed polymeric behavior more consistent with a self-avoid random walk than a compact chain poised to undergo self-assembly (**Supplemental Figure S1d**). This suggests the N-terminus acts as a relatively expanded chain that, in spite of the transient intra-molecular contacts mediated by its aromatic residues, would not be expected to engage extensively in long-range attractive interactions.

### TTDN1 forms condensates in vitro

To evaluate the ability of TTDN1 to form condensates *in vitro*, we generated and purified a TTDN1-mCherry fusion protein (TTDN1-mCherry) from insect cells (**Supplemental Figure S1e**). We initially analyzed condensate formation in the presence of polyethylene glycol (PEG) which readily demonstrated spherical droplet formation of ∼0.5-1 μm size at lower protein concentrations (2.5 µM) (**Figure 1e-f**). These droplets were dynamic, demonstrating both fusion and fission properties during live imaging (**Figure 1g**). Higher concentrations of TTDN1-mCherry permitted condensate formation without PEG (**Figure 1h-i**). To determine the contribution of the N-terminal aromatic residues in droplet formation, we generated an aromatic to alanine TTDN1-mCherry fusion (TTDN1^φ→Ala^-mCherry), targeting all 14 aromatics residues in this domain. This led to a ∼50% reduction in droplet formation (**Figure 1j-k**). Together with the simulations data, these findings indicate that TTDN1 is able to undergo phase separation *in vitro*, and this function relies at least in part on the contribution of intra-molecular contacts mediated by its aromatic residues. While one extrapolation of these data is that TTDN1 drives condensates *in vivo* through homotypic interactions, an alternative interpretation is that the transient aromatic-mediated interactions identified *in silico* and *in vitro* may contribute to heterotypic interactions with other binding partners. As such, we sought to better understand the TTDN1 binding partners to examine if and how these IDR-mediated interactions might contribute to its function.

### TTDN1 interacts directly with DBR1 and promotes RNA lariat processing in cells

To identify the proteomic network with which TTDN1 may interact, we used mass spectrometry analysis of tagged TTDN1 immunopurified from HeLa-S nuclear extracts, as HA-TTDN1 localizes to the nucleus (**Supplemental Figure S2a**). From two independent purifications, the intron lariat debranching enzyme, DBR1, was isolated with TTDN1 (**Supplemental Table 1** and **Supplemental Figure S2b**). We confirmed this interaction by immunoprecipitation of HA-TTDN1, which bound to endogenous DBR1 (**Figure 2a**), as well as reciprocal immunoprecipitation of endogenous DBR1 (**Figure 2b**). To determine whether TTDN1 and DBR1 could interact directly, we immobilized recombinant GST-TTDN1 and tested the ability to bind recombinant His-Flag-DBR1. While GST-TTDN1 was able to pull down His-Flag-DBR1, two negative controls (GST alone and GST-ASCC1) did not (**Figure 2c**).

**Figure 2.**
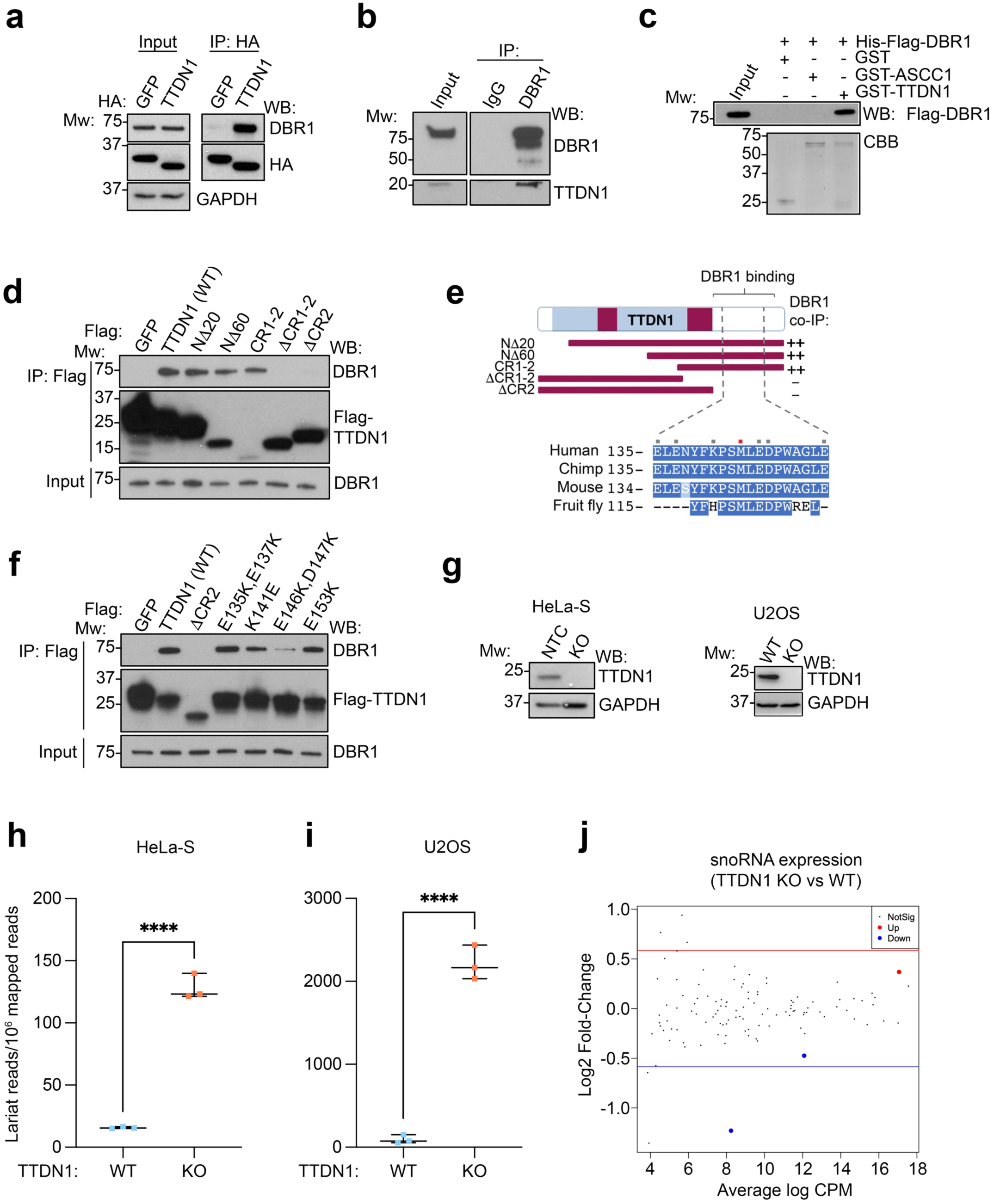
TTDN1 interacts directly with the lariat debranching enzyme DBR1 and promotes RNA debranching in cells. **(a)** HA IP was performed from 293T cells expressing HA-GFP or HA-TTDN1. IP and input material were analyzed by Western blot as shown, with positions of molecular weight (Mw) markers shown on the left. Figure representative of three independent experiments. **(b)** Whole cell extracts from 293T cells were immunoprecipitated with control IgG or DBR1 antibody. IP and input material were analyzed by Western blot as shown. Figure representative of three independent experiments. **(c)** GST, GST-ASCC1, and GST-TTDN1 were immobilized and binding with full-length recombinant His-Flag-DBR1 was tested. Bound and input material were analyzed by Western blot (top) or Coomassie Blue staining (CBB; bottom). Figure representative of three independent experiments. **(d)** Flag IP was performed from 293T cells expressing the indicated Flag proteins. IP and input material were analyzed by Western blot as shown. Figure representative of three independent experiments. **(e)** Schematic of TTDN1 and summary of DBR1 binding analysis. CR1 and CR2 refer to “conserved region 1” and “conserved region 2” (Nakabayashi *et al*., 2005). Bottom shows C-terminal sequence alignment of human TTDN1 with orthologues from other species. Red square indicates site of NP-TTD associated point mutation (Met144→Val). Grey squares indicate residues targeted for mutagenesis. **(f)** Flag IP was performed as in **(d)** using the indicated vectors. IP and input material were analyzed by Western blot as shown. Figure representative of three independent experiments. **(g)** TTDN1 was targeted using CRISPR/Cas9 in HeLa-S and U2OS cells. Individual clones were isolated and analyzed by Western blot using antibodies against TTDN1 and GAPDH. **(h)** and **(i)** RNA-Seq was performed in the indicated cell lines, and stable lariat species were identified and quantified using a branchpoint detection algorithm in HeLa-S **(h)** and U2OS cell lines **(i)**. N = 3 RNA-Seq replicates and lariat analysis per cell line; **** *p* < 0.0001 by unpaired t-test. **(j)** Bland-Altman plot comparing snoRNA expression in control and TTDN1 KO HeLa-S cells. Red and blue lines indicate Log_2_ fold-change of +/-0.585, corresponding to linear fold change of +/-1.5.

To identify the region of TTDN1 required for interaction with DBR1, we performed a deletion analysis of TTDN1. The N-terminus of TTDN1 was dispensable for this interaction, while the CR1-2 domain was both necessary and sufficient for binding DBR1 (**Figure 2d-e**). The M144V patient mutation, located within the CR2 domain, modestly affected the interaction with DBR1 (**Supplemental Figure S2c**). Additional site-directed mutagenesis demonstrated that specific charged residues within this domain, in particular E146 and D147, were important for the DBR1 interaction (**Figure 2e-f, Supplemental Figure S2c**).

Because of the direct interaction between TTDN1 and DBR1, we reasoned that TTDN1 may play a role in lariat processing by DBR1. Therefore, we generated CRISPR/Cas9 clonal knockouts of TTDN1 in HeLa-S and U2OS cells (**Figure 2g**). To quantify lariat processing, we performed RNA-seq at high depth (>200 million reads/sample), then applied a modified version of a previously described branchpoint annotation algorithm (Pineda and Bradley, 2018). Using this approach, we found that loss of TTDN1 increased total lariat abundance ∼8.2 and ∼23.7-fold over controls in HeLa-S and U2OS cells, respectively (**Figure 2h-i**). This was not due to loss of DBR1 protein (**Supplemental Figure S2d** and **Figure 6** below). Our RNA-Seq analysis also revealed that loss of TTDN1 led to aberrant splicing events, which were found to be primarily exon skipping (**Supplemental Figure S2e-f**). A similar phenotype has been previously observed in cells expressing lower levels of DBR1 (Han *et al*., 2017). We corroborated this by depleting DBR1 and performing RNA-Seq to assess splicing event alterations (**Supplemental Figure S2g-h**). Since the processing of small nucleolar RNAs (snoRNAs) also depends on DBR1 (Hirose *et al*., 2003), we performed RNA-seq after small RNA isolation using the TTDN1 KO and control HeLa-S lines to determine misregulated expression of this class of small RNAs. Virtually none of the snoRNAs were significantly altered in TTDN1 KO cells as compared to WT controls, suggesting that introns encoding snoRNAs may not be affected by TTDN1 loss (**Figure 2j**).

### TTDN1 links DBR1 to the intron binding complex

One way in which TTDN1 could be promoting DBR1 activity was to serve as a tether for association with higher-order RNPs. To test this, we performed size exclusion chromatography on nuclear extracts from control and TTDN1 KO cells (**Supplemental Figure S3a**). Most of the endogenous DBR1 from both extracts eluted at a lower molecular weight (∼150 kDa). However, in WT nuclear extract, a small amount of DBR1 co-eluted with earlier fractions, suggesting DBR1 may associate with larger complexes, which appeared lost in TTDN1 KO extracts (**Supplemental Figure S3a**, fractions 26-30). Interactome analysis of DBR1 immunopurified from HeLa-S nuclear extracts revealed that a majority of DBR1-interacting proteins were associated with the spliceosome (**Supplemental Table 2**), consistent with previous reports (Masaki et al., 2015). Enriched among these were all five members of the intron binding complex (IBC), composed of the RNA helicase Aquarius (AQR), XAB2, ISY1, ZNF830, and PPIE), which incorporates upstream of the intron branch point into catalytically active spliceosomes (**Supplemental Figure S3b**) (De et al., 2015; Hirose et al., 2006). Notably, many of these interaction partners appeared reduced or lost in the DBR1 IP-MS performed in parallel from TTDN1 KO cells (**Figure 3a** and **Supplemental Table 3**). IP-Western analysis of tagged DBR1 confirmed interaction with IBC members in WT extract, but strikingly, the IBC interaction with DBR1 was lost in TTDN1 KO cells (**Figure 3b**), indicating that TTDN1 may function to tether DBR1 and the IBC.

**Figure 3.**
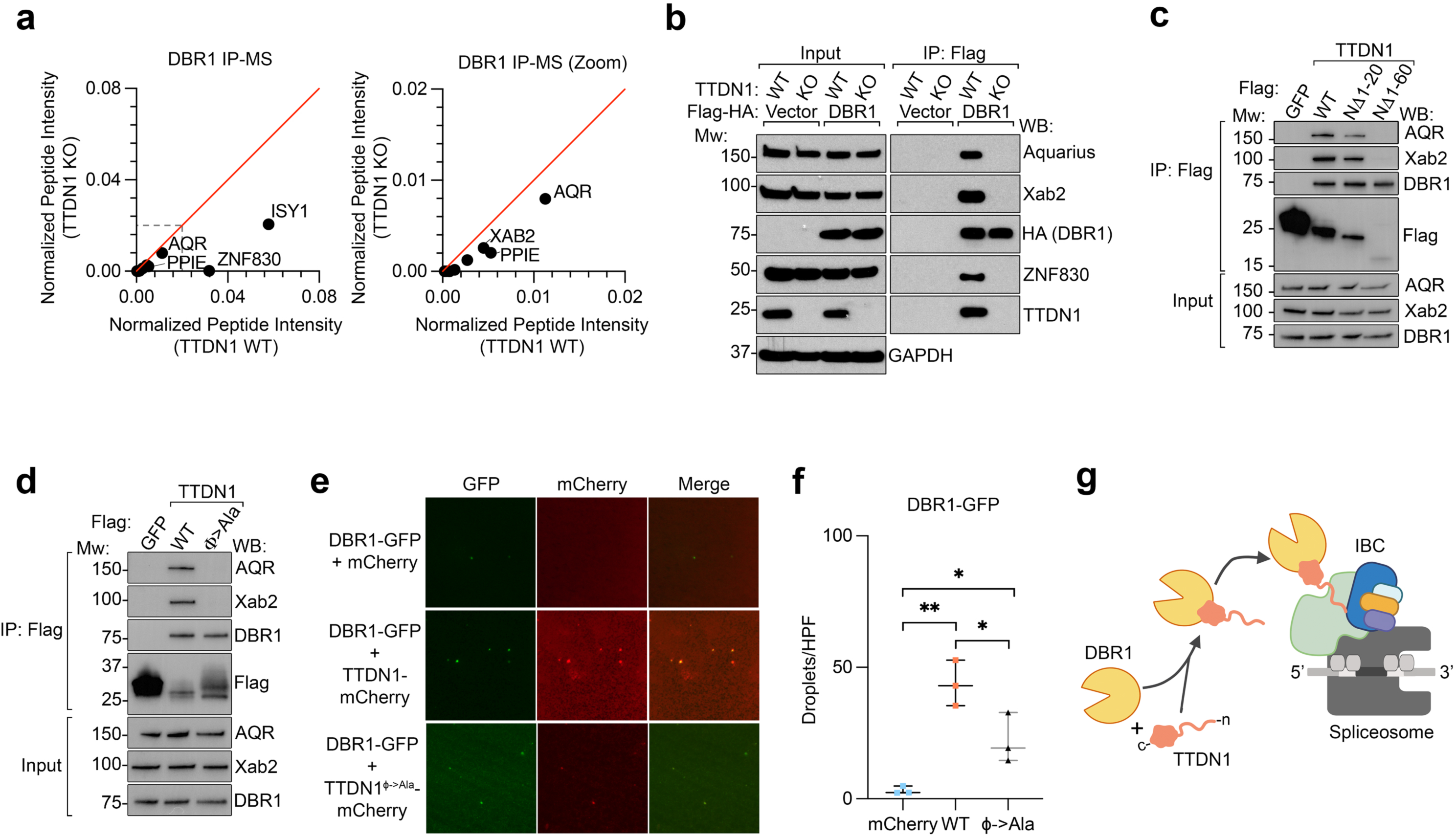
TTDN1 bridges DBR1 to the intron binding complex. **(a)** Peptide plots depicting normalized sum intensities (averaged from two independent experiments) for proteins associated with DBR1 in control or TTDN1 KO HeLa-S cells expressing Flag-HA-DBR1, as determined by mass spectrometry. **(b)** Flag-HA-DBR1 or vector control was expressed in WT or TTDN1 KO HeLa-S cells. Flag immunoprecipitation was performed from nuclear extract and Western blotted as shown. Figure representative of three independent experiments. **(c)** and **(d)** Flag-tagged vectors were transiently expressed and immunoprecipitated from 293T cells, then Western blotted as shown. Each figure representative of three independent experiments. **(e)** Condensate formation of GFP-DBR1 (0.15 µM) in the presence of mCherry, TTDN1-mCherry or TTDN1^φ→Ala^-mCherry (2.5 µM each). Scale bar, 10 µm. **(f)** Quantification of GFP-DBR1 droplet formation from **(e)**. N = 3 independent replicates per condition; * *p* < 0.05, ** *p <* 0.01, n.s., not significant, by unpaired t-test. **(g)** Model for the tethering of DBR1 to the IBC via TTDN1. See text for details.

We reasoned that tagging TTDN1 at the N-terminus with Flag and HA interfered with the ability of TTDN1 to bind the IBC, thus explaining the lack of IBC peptides in our original TTDN1 interactome analysis. Indeed Flag-only tagged TTDN1 repeatedly co-immunoprecipitated XAB2 and AQR, two of the IBC components (**Figure 3c**). While deletion of the first N-terminal 20 amino acids of TTDN1 had a modest effect on this interaction, deletion of 60 amino acids from this region abrogated XAB2/AQR co-immunoprecipitation without impacting DBR1 binding. In addition, the TTDN1^φ→Ala^ condensate-deficient mutant lost its ability to interact with AQR and XAB2 (**Figure 3d**). Thus, the capacity of TTDN1 to form condensates may be linked to its interaction with the IBC.

In this context, we wondered whether condensate formation by TTDN1 might be relevant to its function with DBR1, which is predicted to have a disordered region in its C-terminus (**Supplemental Figure S3c**). We generated and purified a DBR1-GFP fusion protein from insect cells (**Supplemental Figure S3d**) to determine whether DBR1 can incorporate into TTDN1 droplets. Relative to mCherry alone, droplets formed by DBR1-GFP increased significantly (∼13-fold) with TTDN1-mCherry, and substitution of TTDN1^φ→Ala^-mCherry reduced incorporation by approximately 50% (**Figure 3e-f**). Consistent with our results showing the N-terminus of TTDN1 is dispensable for interacting with DBR1, binding of DBR1-GFP to immobilized TTDN1^φ→Ala^-mCherry appeared equal to TTDN1-mCherry (**Supplemental Figure S3e**). We conclude that DBR1 and TTDN1 can co-localize within phase-separated droplets, mediated at least in part by the N-terminal aromatic residues of TTDN1 *in vitro*. Collectively, our data support a model in which the C-terminus of TTDN1 functions to recruit DBR1, while the condensate-forming N-terminus of TTDN1 promotes IBC interaction, bringing DBR1 to nascent RNA lariats (**Figure 3g**).

### Lariat processing defects influence gene expression in a length-dependent manner

How might disruption of this RNP-DBR1 interaction cause TTD? In transcription-coupled repair defective disorders, long genes are thought to accumulate a higher total lesion load, or increased number of lesions, than short genes. Over time, this results in biased misexpression of genes in a length-dependent manner due to RNA Pol II stalling (Vermeij et al., 2016a). While patients with *TTDN1* alterations are repair-proficient, they exhibit severe developmental and neurological phenotypes that are consistent with what is seen in TC-NER defective TTD patients; yet whether or how TTDN1 loss correlates with any of these transcriptional defects is unclear (Faghri *et al*., 2008). Strikingly, we found that the average genomic length of downregulated transcripts in the absence of TTDN1 was significantly longer than the length of unaffected genes in both the U2OS and HeLa-S cell lines (**Figure 4a** and **Supplemental Figure S4a**). A similar inverse correlation was observed when analyzing the number of exons and gene expression changes (**Supplemental Figure S4b-c**). Plotting gene expression change compared to transcript genomic length consistently revealed an inverse relationship between these two parameters in the TTDN1 KO cells relative to controls (**Figure 4b** and **Supplemental Figure S4d**). We confirmed that three long genes (*TLL1*, *BCR*, and *POU6F2*) which were significantly downregulated in our RNA-Seq data sets were similarly reduced by qRT-PCR in the TTDN1 KO cells versus WT controls (**Figure 4c**).

**Figure 4.**
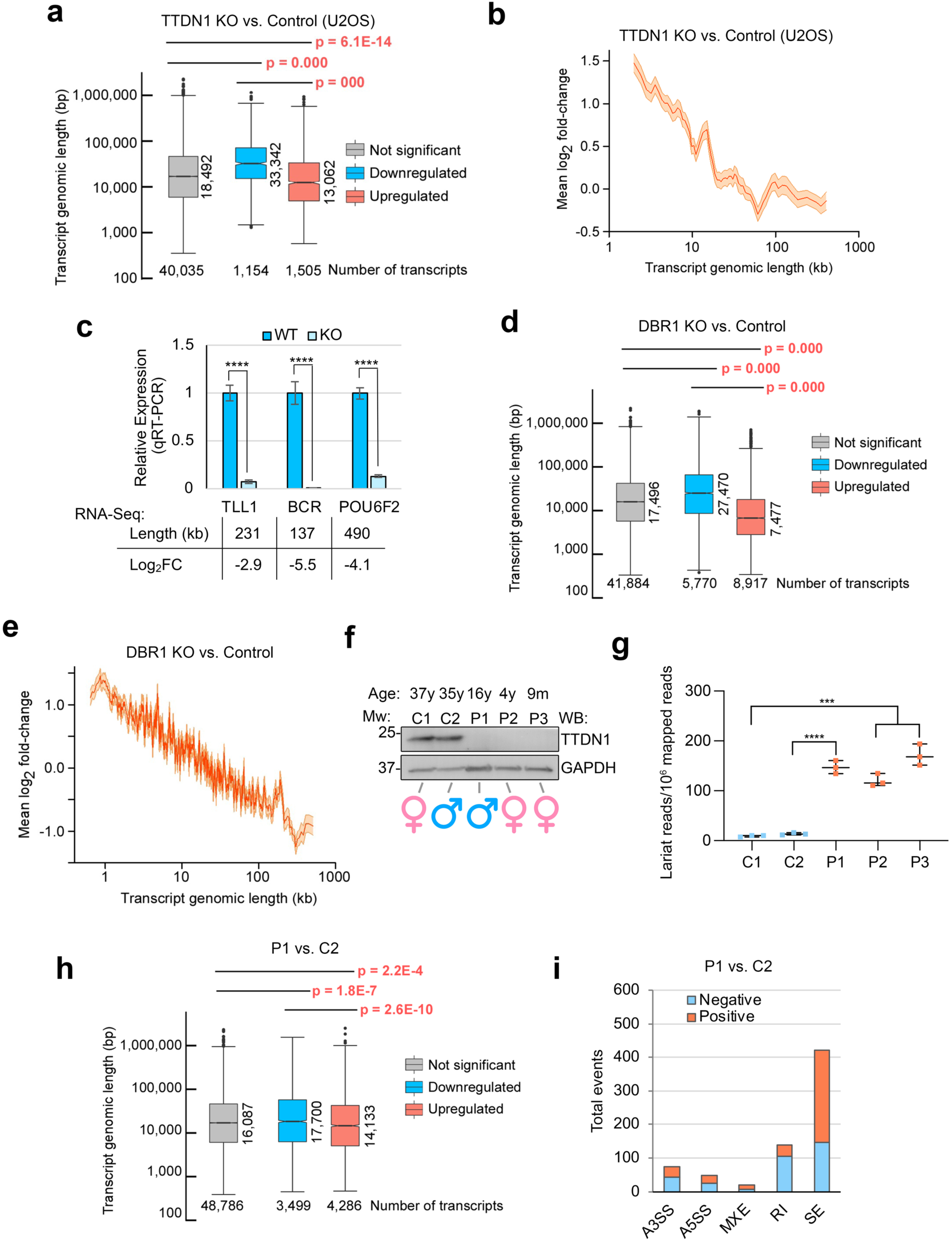
Length-dependent gene expression changes upon loss of *TTDN1* and *DBR1*. **(a)** Box plots of transcript genomic length of DEGs (<-0 log2 fold-change (*Downregulated*), >0 log2 fold-change (*Upregulated*), and not differentially expressed genes (*Not significant*)) from TTDN1 KO and control U2OS cells. *p* values were determined by Wilcoxon rank-sum tests. **(b)** Inverse relationship between transcript genomic length and changes in expression (log2 fold-changes) upon loss of TTDN1 in U2OS cells. **(c)** mRNA levels of indicated genes were assessed by RT-qPCR in U2OS WT and TTDN1 KO cells. Internal expression control was *β*-actin. Genomic transcript length and Log_2_ fold change values from corresponding RNA-Seq results are displayed below graph. Error bars represent standard deviation of data from two independent experiments. **** *p* < 0.0001 by unpaired t-test. **(d)** and **(e)** Analysis as in **(a)** and **(b)**, respectively, was performed using DEGs from RNA-Seq analysis of pooled DBR1 KO and control HeLa-S cells. **(f)** Whole cell lysates from two control (C1 and C2) and three NP-TTD (P1, P2, and P3) patient fibroblast lines with known *TTDN1* mutations were used for Western blot analysis with the indicated antibodies. **(g)** Stable lariat species were identified and quantified using a branchpoint identification algorithm in cell lines from **(f)**. N = 3 RNA-Seq replicates and lariat analysis per cell line; *** *p* < 0.001 **** *p* < 0.0001 by unpaired t-test. **(h)** Analysis as in **(a)** was performed using DEGs from RNA-Seq analysis of P1 and C2 patient cells. **(i)** Alternative 3′ and 5′ splice sites (A3SS, A5SS), mutually exclusive exons (MXE), retained introns (RI), and skipped exons (SE) were quantified using rMATS analysis on n=3 technical replicate RNA-Seq samples from P1 and C2 patient cells.

The effect of DBR1 loss on length-dependent gene expression was even more striking (**Figure 4d-e**), suggesting the downstream effects of aberrant lariat processing in the absence of TTDN1, albeit not as severe as DBR1 loss, are sufficient to result in similar gene expression changes. We then extended this analysis to patient fibroblasts from three siblings with NP-TTD — all homozygous for a two base pair deletion at nucleotides 187-188 in exon 1 of *TTDN1* resulting in an early frameshift (Nakabayashi *et al*., 2005). Consistent with the prediction that these patients are genetically null, we could not detect TTDN1 protein by Western blot in all three patient fibroblasts (**Figure 4f**). Again, using high-depth RNA-seq, we found an increase in RNA lariat accumulation in each NP-TTD fibroblast line, validating increased lariat abundance observed in our engineered KO cells reflects the molecular defect in NP-TTD patients (**Figure 4g**). When assessing gene misexpression in the patient fibroblast RNA-Seq data, we saw more modest alterations with respect to gene length, although shorter genes were consistently upregulated in the NP-TTD patient cells compared to controls (**Figure 4h** and **Supplemental Figure S4e, S4g**). This could be an artifact of mismatched patient age, or that the control fibroblasts were from unrelated individuals. Notably, these patient fibroblasts had similar altered splicing patterns seen in our TTDN1 KO cell lines (**Figure 4i** and **Supplemental Figure S4f, S4h, S2e-f**), suggesting a similar defect in mRNA processing in these patient cells.

### Loss of *Ttdn1* in mice recapitulates RNA processing defects and NP-TTD pathology

Our *in vitro* findings support a model in which lariat accumulation leads to splicing disruption and defects in length-dependent gene expression. We next asked how loss of TTDN1 *in vivo* could lead to downstream consequences on development and neurological function. While there are multiple NER-deficient mouse models (Vermeij et al., 2016b), to date none of these models recapitulate phenotypes resulting from repair-proficient TTD. As such, we created a mouse model for the common TTDN1 allele found in NP-TTD patients (*TTDN1^M144V/M144V^*) (Nakabayashi *et al*., 2005). Using CRISPR/Cas9, we produced the corresponding homozygous mouse (*Ttdn1^M143V/M143V^*), as well as a large end-joining mediated deletion that resulted in a premature stop codon, leading to truncation of the *Ttdn1* transcript and lack of detectable protein expression (hereon referred to as *Ttdn1^Δ/Δ^*; **Supplemental Figure S5a-b**). We saw significant defects in weight gain over time in both female and male *Ttdn1^Δ/Δ^* mice (**Figure 5a** and **Supplemental Figure S5c**), consistent with what is seen in most NP-TTD patients (Faghri *et al*., 2008). However, we failed to see this phenotype in the *Ttdn1^M143V/M143V^* mice compared to littermate controls (**Supplemental Figure S5d**). Indeed, patients with M144V substitution appear to have a less severe phenotype than other TTDN1 alleles (Nakabayashi *et al*., 2005), likely reflecting the modest reduction in its ability to interact with DBR1 (**Supplemental Figure S2c**). From heterozygous matings, *Ttdn1^Δ/Δ^* mice were born at less than expected Mendelian ratios, likely reflecting a modest reduction in overall fitness (**Supplemental Figure S5e**). A hallmark phenotype in TTD is the presence of sparse, sulfur-deficient hair, and while this was not overtly apparent in younger *Ttdn1^Δ/Δ^* mice (**Figure 5b**, left), we observed a significant decrease in cysteic acid content as a percentage of total hair protein in these mice (**Figure 5c**). This reduction is on par with decreased levels of total cysteine content observed in hair samples from both TTD patients and a previously-characterized NER-defective TTD mouse model (de Boer et al., 1998). While most TC-NER deficient mice die prematurely due to symptoms of premature aging, *Ttdn1^Δ/Δ^* mice did not appear to grossly deteriorate or die early, as aged *Ttdn1^Δ/Δ^* mice are still alive beyond 15 months, reflecting the lack of progeroid phenotypes in NP-TTD patients. However, the presence of sparse hair became more apparent in the aged *Ttdn1^Δ/Δ^* mice (**Figure 5b**, right), while defects in weight gain as compared to littermate controls were maintained over time (**Supplemental Figure S5f**).

**Figure 5.**
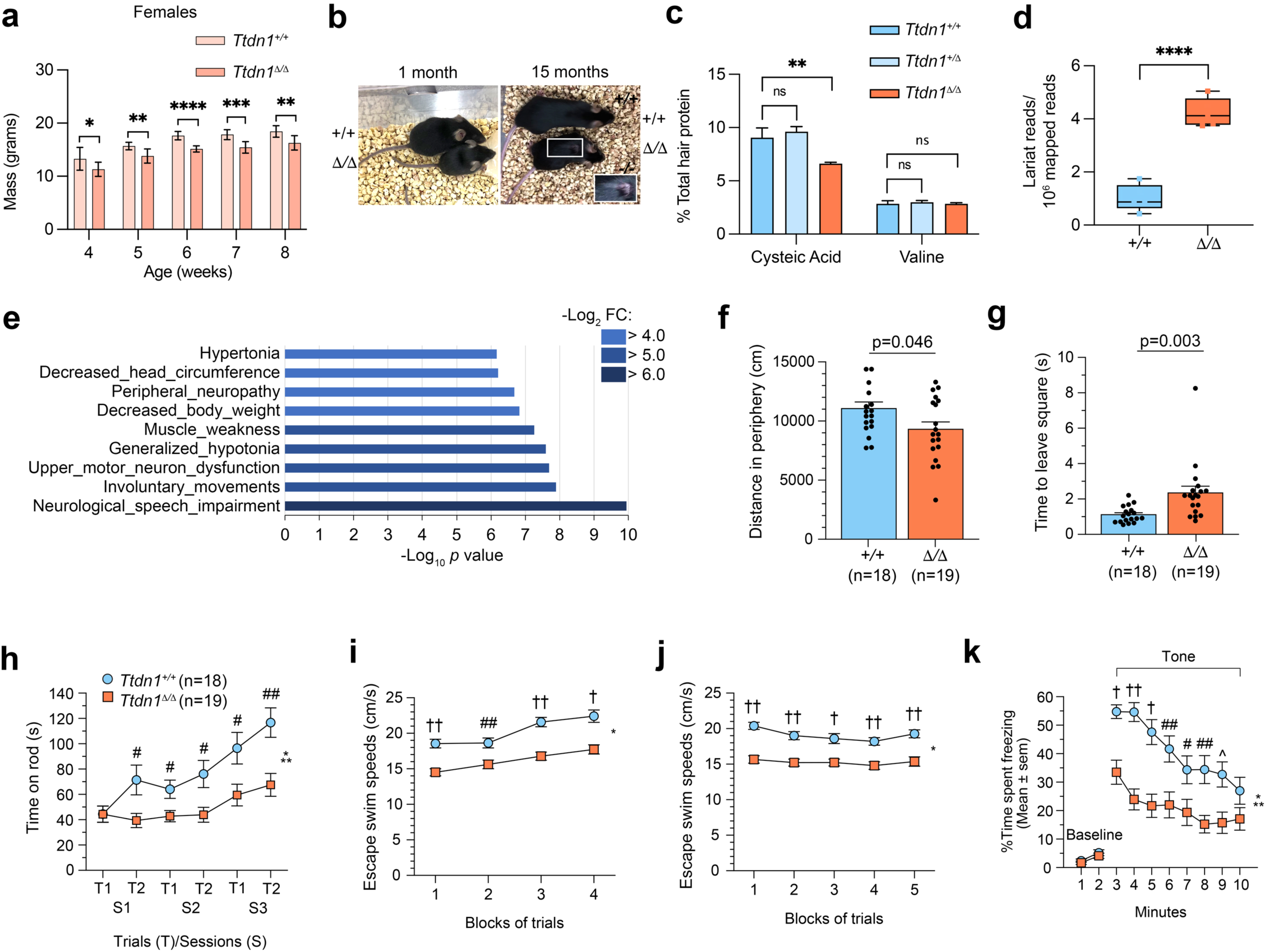
*Ttdn1^Δ/Δ^* mice recapitulate molecular and pathological phenotypes of NP-TTD. **(a)** Weights of female littermate mice were determined at the indicated age. N = 5 mice per genotype. * p < 0.05, ** p<0.01, *** p< 0.001, ****p<0.0001 by unpaired t-test. **(b)** Photograph showing female littermate *Ttdn1^+/+^* and *Ttdn1^Δ/Δ^* mice at ages 1 month (left) and 15 months (right). Note sparse hair is more apparent in aged *Ttdn1^Δ/Δ^* mouse. **(c)** Amino acid analysis of total hair protein from littermate mice. N = 5 per genotype, ** *p* < 0.01, by unpaired t-test. **(d)** RNA was extracted from the cortex of 8 week old littermate *Ttdn1^+/+^* and *Ttdn1^Δ/Δ^* mice. Stable lariat species were identified and quantified using a branchpoint identification algorithm. N = 5 mice per genotype. **** p <0.0001 by unpaired t-test. **(e)** GSEA of the top 9 pathways significantly differentially regulated in the cortex of 8 week old *Ttdn1^Δ/Δ^* mice based on MSigDb Human Phenotype Ontology gene sets. **(f)** An ANOVA conducted on the data pertaining to distance traveled in the peripheral zone of the test field showed that the *Ttdn1^Δ/Δ^* mice traveled a significantly shorter distance than controls in this area [F(1,33)=4.32, p=0.046]. **(g)** An ANOVA performed the walking initiation test (combined cohorts) data yielded a significant genotype effect [F(1,33)=15.07, p=0.003], showing that the *Ttdn1^Δ/Δ^* mice took significantly longer to move out of a small circumscribed area. **(h)** An rmANOVA conducted on the data from the accelerating rotarod trials produced a significant genotype effect [F(1,33)=11.74, *p=0.002,], and genotype x trials interaction [F(1,33)=6.67, indicating that the *Ttdn1^Δ/Δ^* mice spent significantly less time on the rotarod for some of the trials **p=0.014]. #p<0.025; ##p<0.003 **(i)** and **(j)** Significant genotype effects were found following rmANOVAs conducted on the swimming speed data from the cued and place trials conducted in the Morris water maze, [genotype effects: F(1,33)=7.87, * p <0.00005; and [F(1,33)=33.86, *p<0.00005, respectively], indicating that *Ttdn1^Δ/Δ^* mice swam significantly more slowly than control mice. A significant sex effect was found during the cued trials, [Sex effects on cued trials: F(1,33)=7.87, p=0.008], but the genotype x sex interaction was not significant. ##p<0.015; †p<0.0005; ††p<0.00005 **(k)** An rmANOVA conducted on the auditory cue data from the conditioned fear test (day 3) resulted in a significant genotype effect, [F(1,33)=20.69, *p=0.001], and a significant genotype x minutes interaction, [F(7,231)=2.33, **p=0.033, Huyhn-Feldt (H-F) adjusted p], showing that the *Ttdn1^Δ/Δ^* mice exhibited significantly reduced freezing levels for certain times during the test #p<0.05, ^p<0.010, ##p<0.00625 (Bonferroni corrected level); †p<0.0005; ††p<0.00005.

To determine whether the *Ttdn1^Δ/Δ^* mice recapitulated the RNA processing defects we saw in our cell lines and patient fibroblasts, we performed RNA-seq from the cortex of *Ttdn1^Δ/Δ^* and wildtype littermate controls. In choosing a tissue to sequence, we considered that the cortex is frequently connected with neurodevelopmental disorders (Chen et al., 2015; Gabel et al., 2015). RNA lariat abundance in these samples was increased ∼4.1-fold compared to WT samples, as were alternative splicing events (**Figure 5d** and **Supplemental Figure S5g**). Gene set enrichment analysis of pathways significantly differentially dysregulated in the cortex of *Ttdn1^Δ/Δ^* revealed molecular signatures associated with neurological and developmental defects in humans (**Figure 5e**). Male *Ttdn1^Δ/Δ^* brains were smaller than controls, whereas female brains were not significantly different (**Supplemental Figure S5h).** Together, these data suggest a connection between molecular defects in the *Ttdn1^Δ/Δ^* mice and the pathology seen in NP-TTD.

### Behavioral Assessment of Ttdn1^Δ/Δ^ Mice

To further characterize their phenotypes, *Ttdn1^Δ/Δ^* mice and *Ttdn1^+/+^*controls from two different cohorts were evaluated on several behavioral tasks including a 1-hour locomotor activity test, a marble-burying test, and a battery of sensorimotor measures (walking initiation, ledge platform; pole, 60° and 90° inclined screens, inverted screens). All of the mice were assessed on the same behavioral testing procedures and test sequence. Except for a modest, but significant reduction in distance traveled in the peripheral zone, *Ttdn1^Δ/Δ^* and control mice were not significantly different on variables related to locomotor activity (**Figure 5f** and **Supplemental Figures S6a-c**). Many NP-TTD patients have autistic-like behaviors, thus mice were also evaluated on the marble-burying test, an often-used measure for assessing models of autism (Heller et al., 2015). However, *Ttdn1^Δ/Δ^* and control groups did not differ in terms of compulsive digging, as measured by the number of marbles buried during the test (**Supplemental Figure S6d**). Out of the total seven measures within the sensorimotor battery, a significant genotype effect was found for only the walking initiation test in terms of the time taken to move out of a small circumscribed area, suggesting fear of moving in an open, novel environment or a slowed motor response in the *Ttdn1^Δ/Δ^* mice (**Figure 5g**, **Supplemental Table 3**). Fine-motor coordination was assessed using the rotarod test. Significant performance deficits were found in the *Ttdn1^Δ/Δ^* mice for the time they were able to remain on the accelerating rotarod (**Figure 5h**), but not for the stationary or constant speed components of the rotarod procedure (**Supplemental Figures S6e-f**). Together, these data suggest that while loss of *Ttdn1* contributes to defects in fine-motor coordination, several basic sensorimotor functions are largely unaffected.

Spatial learning and memory capabilities were assessed next in the mice using the Morris Water Maze (MWM), followed by an evaluation of associative memory performance using a Pavlovian fear conditioning procedure. While there were no significant deficits in spatial learning and memory in the *Ttdn1^Δ/Δ^* mice (**Supplemental Figures S6g-j**), we observed significantly reduced swimming speeds in the *Ttdn1^Δ/Δ^* mice during the cued and place trials (**Figure 5i-j**), suggesting impaired coordination and/or motivational disturbances. Analysis of the conditioned fear data showed that the *Ttdn1^Δ/Δ^* mice exhibited significantly reduced freezing levels on the auditory cue component (day 3) of the procedure (**Figure 5k and Supplemental Figures S6k-l**). Importantly, no significant differences between knockout and control mice were found in freezing levels during the following: baseline or tone-shock training (day 1); the contextual fear test (day 2); altered context baseline (day 3); or shock sensitivity. Moreover, no significant effects were observed on measures of acoustic startle or pre-pulse inhibition (only tested in cohort 2; **Supplemental Figures S6m-o**). We conclude that the deficit in auditory cue conditioning exhibited by *Ttdn1^Δ/Δ^* mice is a selective cognitive impairment not likely due to deafness or extreme auditory deficits. Altogether, our results suggest that the *Ttdn1^Δ/Δ^* mice likely have impaired fine motor coordination and/or motivational disturbances, as well as specific fear (auditory cue) conditioning deficits.

### U2 snRNP inhibition mirrors TTDN1/DBR1 loss in altering length-dependent gene expression

Determining a unified basis for how NP-TTD shares molecular pathology with photosensitive TTD has remained obscure due to the genetic heterogeneity of the disease. Mutations in TF_II_Eβ (Theil et al., 2017), aminoacyl-tRNA synthetases (Botta et al., 2021; Kuo et al., 2019; Theil et al., 2019), and the splicing protein RNF113A (Corbett *et al*., 2015; Mendelsohn *et al*., 2020; Tessarech *et al*., 2020) have all been linked to NP-TTD. We reasoned that the molecular defect between the latter and TTDN1 may both be explained by a common spliceosomal defect; indeed, RNF113A joins the activated spliceosome just prior to the first transesterification step in splicing (Haselbach et al., 2018), and alterations in RNA Pol II elongation are seen upon loss of the ASCC complex, which is recruited downstream of RNF113A (Brickner *et al*., 2017; Williamson et al., 2017). Therefore, we reasoned that inhibiting the spliceosome upstream of TTDN1 and DBR1 may result in similar length-dependent gene expression changes. We tested the effect of the early-stage spliceosome inhibitor pladienolide-B (Pla-B), which inhibits the U2-associated SF3B complex (Cretu et al., 2018), on gene expression and RNA processing in WT and TTDN1 KO cells using RNA-seq. We first assessed RNA lariat accumulation and found that Pla-B treatment does not have a significant effect on lariat levels at steady state (**Figure 6a**). However, Pla-B treatment leads to a similar length-dependent gene expression change as TTDN1/DBR1 loss, increasing the expression of shorter genes and negatively affecting longer ones (**Figure 6b** and **Supplemental Figure S7a**). This effect of Pla-B was also observed in TTDN1 KO cells (**Supplemental Figure S7b-c**), although we noticed a significantly stronger impact of Pla-B on WT cells versus those deficient for TTDN1 (i.e., a greater number of transcripts were affected by Pla-B in WT cells) (**Figure 6c-d**). Indeed, there was a significant overlap between genes significantly downregulated in TTDN1 KO cells versus Pla-B treated WT cells (**Figure 6e**), which indicates that SF3B inhibition is at least partially epistatic with TTDN1 loss. These data suggest that reduced spliceosomal function, whether early or late in the spliceosome cycle, may lead to a common deficit in length-dependent gene expression.

**Figure 6.**
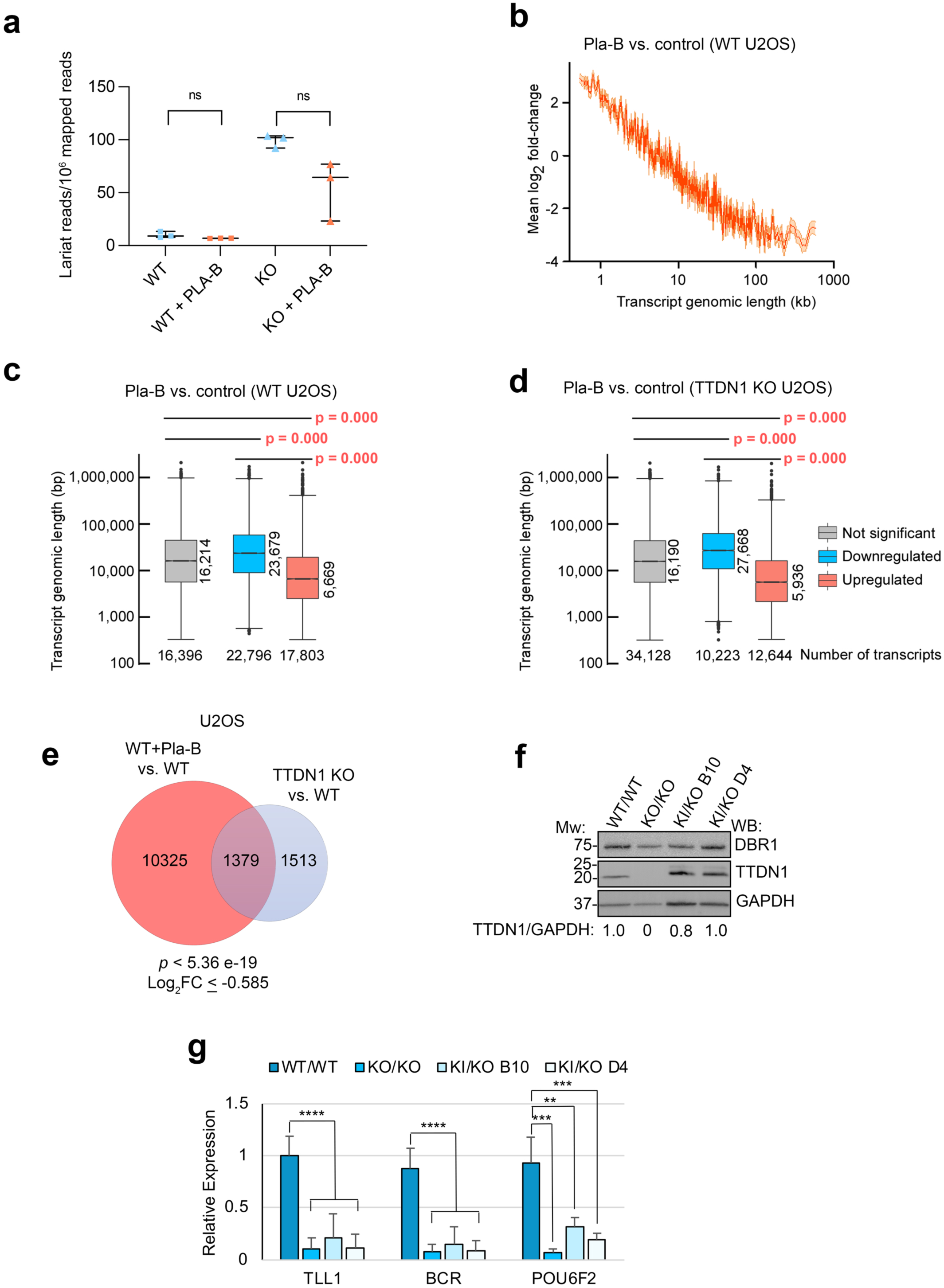
TTDN1 and spliceosome function in length-dependent gene expression. **(a)** RNA-seq was performed in the indicated U2OS cell lines, and stable lariat species were identified and quantified using a branchpoint detection algorithm in the presence or absence of Pladienolide B (Pla-B). N = 3 RNA-Seq replicates and lariat analysis per cell line; ns, not significant by unpaired t-test. **(b)** Inverse relationship between transcript genomic length and changes in expression (log2 fold-changes) in WT U2OS cells upon spliceosome inhibition with Pla-B. **(c)** and **(d)** Box plots of transcript genomic length of DEGs (<-0.585 log2 fold-change (*Downregulated*), >0.585 log2 fold-change (*Upregulated*), and not differentially expressed genes (*Not significant*)) from control and TTDN1 KO U2OS cells in the presence or absence of Pla-B. *p* values were determined by Wilcoxon rank-sum tests. **(e)** Venn diagram showing the number of overlapping DEGs <-0.585 log2 fold-change between WT U2OS cells treated with Pla-B and TTDN1 KO U2OS cells. *p* value was determined by one-sided Fisher’s exact test. **(f)** Protein levels in *TTDN1^φ→Ala/Δ^* clonal U2OS cell lines were compared to WT and TTDN1 KO counterparts by Western blot using antibodies as shown. Normalized fold change in TTDN1 band intensity relative to GAPDH is shown below blot. **(g)** mRNA levels of indicated genes were assessed by qRT-PCR in the indicated clonal U2OS cell lines, with β-actin as the normalization control. ** *p* < 0.01, *** *p* < 0.001, **** *p* < 0.0001 by unpaired t-test.

### The TTDN1 IDR is critical for long gene expression

Because TTDN1 forms condensates *in vitro* through its N-terminal IDR, and appears to promote DBR1 condensate formation and function, we wished to test whether this capacity of TTDN1 was important for its role in maintaining long gene expression. We targeted the human *TTDN1* locus in U2OS cells using CRISPR/Cas9 and substituted its entire exon 1 with the N-terminal aromatics mutant (TTDN1^φ→Ala^, see **Figure 1**). Although we were only able to obtain hemizygous *TTDN1^φ→Ala/Δ^* clones, TTDN1 protein levels in two independent mutant clones were similar to the WT counterpart (**Figure 6f**). We evaluated expression of the same three long genes from **Figure 4c** in these cells using qRT-PCR. Strikingly, the significant reduction in expression of these genes in TTDN1 KO cells was mirrored in the *TTDN1^φ→Ala/Δ^* clones (**Figure 6f**). We should note that reduction of TTDN1 protein levels to <50% of WT levels resulted in very modest gene expression changes (**Supplemental Figure S7d-e**), arguing against a simple dosage effect in our *TTDN1^φ→Ala/Δ^* clones. Together, these results suggest that the aromatic residues in the TTDN1 IDR are important for its function in mediating long gene expression, possibly in the context of recruitment to and/or formation of TTDN1-associated condensates.

### A DBR1^TTDN1-NTD^ fusion rescues IBC interaction, lariat processing, and length-dependent gene expression

Despite the above experimental results, our data do not sufficiently prove that the function of the TTDN1 IDR in relation to DBR1 and the DBR1-IBC interaction is critical for lariat processing and length-dependent gene expression. To determine whether restoring DBR1 association with the IBC would be sufficient to rescue these phenotypes, we generated a fusion of the N-terminal IDR of TTDN1 with the C-terminus of full-length DBR1 (DBR1^TTDN1-NTD^) (**Figure 7a**). We reasoned that this would bypass the requirement for the TTDN1 C-terminal domain in bridging DBR1 and the IBC. This fusion protein was indeed capable of interacting with IBC components in TTDN1 KO cells (**Figure 7b**). Strikingly, RNA-seq analysis revealed that it also rescued lariat levels in TTDN1 KO cells to that of WT controls (**Figure 7c**). Finally, we profiled gene expression in TTDN1 KO cells expressing DBR1^TTDN1-NTD^ and found that relative to control TTDN1 KO cells, expression of the fusion protein rescued gene length phenotypes to near-WT levels (**Figures 7d-g** and **Supplemental Figures S7f-g**). Together, these data strongly support the idea that a primary function of TTDN1 is to serve as a molecular tether between DBR1 and the IBC in order to promote efficient lariat processing, which in turn affects length-dependent gene expression.

**Figure 7.**
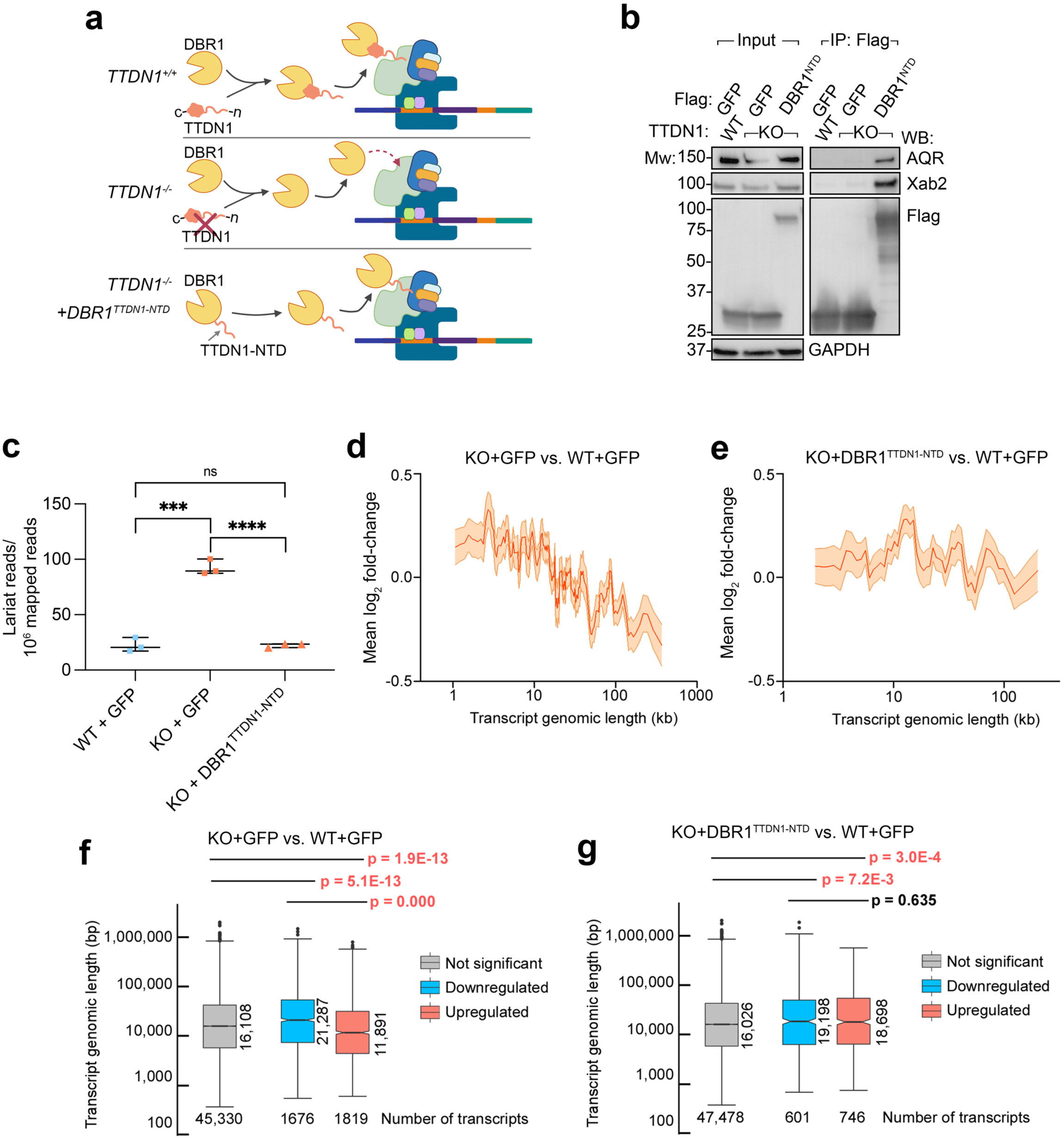
Tethering DBR1 to the intron binding complex rescues TTDN1 deficiency. **(a)** A fusion of DBR1 to the intrinsically disordered domain of TTDN1 is predicted to recruit DBR1 to intron binding complex, bypassing the requirement for TTDN1. **(b)** The indicated Flag vectors were stably expressed in WT or TTDN1 KO HeLa-S cells. Following Flag immunoprecipitation, input and IP material was Western blotted as shown. Figure representative of three independent experiments. **(c)** Stable lariat species were identified and quantified using a branchpoint identification algorithm in cell lines from **(b)**. N = 3 RNA-Seq replicates and lariat analysis per cell line; *** *p* < 0.001 **** *p* < 0.0001 by unpaired t-test. **(d)** and **(e)** Relationship between transcript genomic length and changes in expression (log2 fold-changes) in the indicated HeLa-S cell lines. **(f-g)** Box plots of transcript genomic length of DEGs (<-0.585 log2 fold-change (*Downregulated*), >0.585 log2 fold-change (*Upregulated*), and not differentially expressed genes (*Not significant*)) from **(b)**.

## Discussion

Here we identify TTDN1 as a novel molecule involved in RNA processing, and delineate the connections between aberrant lariat processing upon its loss to gene expression defects in relation to NP-TTD. Notably, we find that the highly conserved C-terminus of TTDN1 functions as an anchor in binding DBR1, whereas aromatic residues within the N-terminal IDR mediate interactions with the IBC. In the absence of TTDN1, we find that the association of DBR1 with active splicing complexes is lost, and that the resulting lariat accumulation consistently coincides with splicing disruption and defects in gene expression across models for TTDN1 deficiency. We observe reduced expression of long genes in particular, which may be a consequence of high intronic burden. In the absence of efficient intron lariat processing, a higher localized requirement for intron lariat debranching at long genes may interfere with splicing, thus inhibiting Pol II elongation. Thus, disruption of early as well as late stages of splicing may significantly alter length-dependent gene expression, suggesting that multiple routes of splicing disruption can have similar consequences for transcriptional outcomes. Our mouse model for NP-TTD phenocopies the hallmark patient hair phenotype, and recapitulates aspects of the developmental and neurological defects seen in patients with TTDN1 deficiency. That expression of a DBR1^TTDN1-NTD^ fusion in TTDN1 KO cells rescues defects associated with TTDN1 loss strengthens the idea that increased coordination between DBR1 and the IBC promotes nascent lariat debranching by DBR1, promoting proper splicing and gene expression.

Photosensitive TTD arises from defects in the transcription-coupled NER pathway. When a bulky adduct, often a result of UV irradiation, is encountered within a transcriptionally active gene, elongating RNA Pol II cannot bypass the lesion and stalls (Egly and Coin, 2011). This triggers the sequential recruitment of various transcription-coupled repair (TCR) proteins that ultimately target TF_II_H to the lesion (Gregersen and Svejstrup, 2018; van der Weegen et al., 2020). The combined action of both the helicase and translocase subunits of TF_II_H, XPD and XPB, contribute to the eventual displacement of the stalled RNA Pol II (Kokic et al., 2019; Sarker et al., 2005). Photosensitivity in TTD cases results from mutations within genes encoding various subunits of TFIIH, affecting UV damage repair, as well as the transcriptional response during such damage. While TF_II_H has important DNA repair functions, it is also associated with multiple stages of transcription, including transcription initiation and re-initiation after Pol II pausing (Henderson et al., 2017; Yudkovsky et al., 2000). This dual function is thought to be responsible for the compound phenotypes seen in TTD patients, specifically as related to tissue-specific transcriptional impairments (Bergmann and Egly, 2001; Dubaele et al., 2003).

TTDN1-mutated patients are non-photosensitive and NER proficient (Botta *et al*., 2007; Heller *et al*., 2015), as are patients with mutations in TF_II_Eβ (Kuschal *et al*., 2016; Theil *et al*., 2017), in the only X-linked TTD-associated gene, RNF113A (Corbett *et al*., 2015; Tessarech *et al*., 2020), and in genes encoding aminoacyl-tRNA synthetases (aaRS) (Botta *et al*., 2021; Kuo *et al*., 2019; Theil *et al*., 2019). However, non-photosensitive cases retain the severe neurological and developmental phenotypes seen in photosensitive patients. While TF_II_Eβ has roles in transcription initiation, RNF113A functions in RNA splicing and recruitment of the ASCC complex, which in turn may affect nascent transcription (Tsao *et al*., 2021; Williamson *et al*., 2017). Although seemingly diverse, this heterogeneity may at least partially converge on gene expression, which has led to the hypothesis that the hallmark features of TTD are a consequence of disturbed gene expression and protein instability, although these explanations do not satisfy the seemingly specific phenotypes in TTD, such as brittle, sulfur deficient hair. (Botta *et al*., 2021). Here, we show that TTDN1 affects the lariat processing activity of DBR1 *in vivo*, which suggests NP-TTD cases can arise from indirect consequences of aberrant splicing culminating in altered nascent transcription and aberrant length-dependent gene expression.

Intron lariat formation begins during two transesterification reactions downstream of spliceosome assembly and catalytic activation. During the first transesterification step, the phosphodiester bond of the 5’ splice site undergoes nucleophilic attack by the 2’OH group of a bulged branch adenosine. This releases the 5’ exon and results in a 2’-5’ phosphodiester linkage between the 5’ splice site and branch adenosine. The second transesterification step results in exon ligation and intron lariat release after the 3’OH of the released 5’ exon attacks the phosphodiester bond of the 3’ splice site (Konarska et al., 1985; Ruskin et al., 1984). The released lariat intron is contained within the Intron Large complex (ILC; (Yoshimoto et al., 2009)), containing U2, U5, and U6 snRNPs, along with several splicing factors. The ATP-dependent DExH box RNA helicase hPrp43 is subsequently recruited and disassembles the ILC to allow DBR1 debranching activity to linearize the intron lariat (Yoshimoto *et al*., 2009). In *Saccharomyces cerevisiae, DBR1* is not essential for viability, and despite increased levels of lariat intron RNAs, there is little growth defect (Chapman and Boeke, 1991). In contrast, the *Schizosaccharomyces pombe dbr1*^null^ mutant has overt growth defects that coincide with cellular elongation and increased lariat intron RNAs (Nam et al., 1997). That introns occupy 95% of protein-coding transcripts in humans may explain why DBR1 is essential (Findlay et al., 2014; International Human Genome Sequencing, 2004; Lander et al., 2001). Thus, while DBR1 is evolutionarily well-conserved, the increasing consequences of DBR1 deficiency correlate with increased intronic burden, and therefore a higher demand for intron turnover.

Yet, it is likely that toxicity associated with DBR1 deficiency is multi-faceted. DBR1 deficiency in humans coincides with intron lariat accumulation concurrent with retention of snRNPs in IL complexes (Han *et al*., 2017). The limited recycling of snRNPs is thought to reduce interactions between active splicing complexes and introns containing weak U2-binding sites, resulting in exon skipping, an alternative splicing (AS) event in which exons following introns with weak branch sites are omitted from mature transcripts. In addition, the role of DBR1 in debranching intron lariats has been connected with the release and processing of various non-canonical regulatory microRNAs (Okamura *et al*., 2007; Ruby et al., 2007) and intronic small nucleolar RNAs (snoRNAs), which function in modification of ribosomal RNAs (Hirose *et al*., 2003). snoRNA deficiency is associated with the production of unmodified rRNAs and reduced ribosome processivity. However, while loss of TTDN1 does not appear to affect all functions of DBR1, such as snoRNA levels, the degree to which lariat accumulation occurs in the absence of TTDN1 is less than that of DBR1. This correlates with the degree of gene expression changes seen upon loss of TTDN1 versus DBR1.

Why is TTDN1 required to tether DBR1 and the IBC? The TTDN1 N-terminus has several putative phosphorylation sites, and previous studies indicate that TTDN1 interacts with the kinases Polo-like Kinase 1 (Plk1) and Cdk1 in a phosphorylation-dependent manner during mitosis (Zhang et al., 2007). Thus, the state of TTDN1 phosphorylation may regulate its interactions with DBR1 or the IBC. However, while TTDN1 phosphorylation is restricted to M phase, TTDN1 expression is maintained throughout the cell cycle (Zhang *et al*., 2007). While DBR1 and IBC members have homologs in *S. pombe*, TTDN1 conservation is restricted to metazoans. Increased intronic burden is a well-characterized phenomenon that coincides with the evolution of higher eukaryotes. Therefore, coordinating lariat processing and intron splicing by TTDN1 may represent a key adaptive strategy to regulate genomic demands seen in multicellular eukaryotes (Frumkin et al., 2019; Verta and Jacobs, 2022).

## Limitations of study

While we provide evidence that TTDN1 forms phase separated droplets *in vitro*, how this promotes DBR1 enzymatic activity is unclear. In addition, the mechanistic basis of how loss of either TTDN1 or DBR1 leads to transcriptional alterations in a length-dependent manner is not fully elucidated. In a related question, whether lariat accumulation is itself potentially toxic to the splicing or transcriptional machineries is unknown. Future studies will test implications of the role of TTDN1 to tether DBR1 and the IBC and further elucidate these mechanistic questions and their connections to TTD pathology.

## Acknowledgments

We thank Zhongsheng You, Hani Zaher, Harrison Gabel, Diana Christian, and members of the Mosammaparast lab for suggestions and assistance during the course of these studies. We thank John Schultz at the UC Davis molecular structure facility for the amino acid analysis, Ross Tomaino at the Harvard Taplin Mass Spectrometry Core for proteomics studies, the Genome Technology Access Center (GTAC), and the Genome Engineering and iPSC (GEiC) Center at Washington University. We acknowledge the Extreme Science and Engineering Discovery Environment (XSEDE, PSC allocations TG-BIO160040 and TG-MCB170053), which is supported by NSF grant ACI-1548562, and the Texas Advanced Computing Center (TACC, http://www.tacc.utexas.edu) at The University of Texas at Austin for providing HPC resources. This work was supported by the NIH (F31 CA0254143 to B.A.T., R35 CA220430 to J.A.T., R01 GM 105681 and R01 GM127472 to W.G.F., R01 CA227001 to N.M., and P01 CA092584 to J.A.T. and N.M.), the Edward P. Evans Foundation (M.J.W.), the Foundation for Barnes-Jewish Hospital Cancer Frontier Fund (M.J.W.), an American Cancer Society Research Scholar award (RSG-18-156-01-DMC to N.M.), the Barnard Foundation (N.M.), Centene Corporation (N.M.), and the Siteman Cancer Center (N.M. and M.J.W.). J.A.T. is also supported by Cancer Prevention Research Institute of Texas (CPRIT) grant RP180813 and the Robert A. Welch Chemistry Chair. A.S.H. is supported by a Longer Life Foundation grant (a collaboration between RGA and Washington University). Partial support for the mouse behavioral studies was provided by the Intellectual and Developmental Disabilities Research Center at the Washington University School of Medicine (P50 HD10342 to D.W.).

## Author Contributions

B.A.T., T.R., N.S., N.E.C., F.M., R.A.S., J.R.B., D.M., M.S.T., and N.M. carried out cellular, biochemical, and animal experiments. B.A.T., L.B., A.B., and S.N.S. carried out bioinformatic analysis. D.F.W. analyzed mouse behavioral studies. A.S.H. performed computer simulations. M.J.W. supervised S.N.S. J.A.T. supervised A.B., W.G.F. supervised L.B. and N.E.C. N.M. supervised the project and wrote the manuscript with B.A.T., with input from all other authors.

## STAR Methods

### RESOURCE AVAILABILITY

#### Lead Contact

Further information and requests for resources and reagents should be directed to and will be fulfilled by the Lead Contact, Nima Mosammaparast.

#### Materials Availability

All reagents generated in this study are available from the Lead Contact without restriction.

#### Data and Code availability

- RNA-seq data will be deposited at GEO and will be made publicly available prior to publication, or upon request.
- Links for original code are provided in the Key Resources Table.
- Any additional information required to reanalyze the data reported in this paper is available from the Lead Contact upon request.

### EXPERIMENTAL MODEL AND SUBJECT DETAILS

#### Cell culture

Human cell lines (293T, HeLa-S, and U2OS; all originally from ATCC) were cultured in Dulbecco’s modified eagle medium (Invitrogen), supplemented with 10% fetal bovine serum (FBS; Sigma), 100 U/ml of penicillin-streptomycin (Gibco) at 37°C and 5% CO_2_. Unaffected control and NP-TTD patient fibroblast cell lines (obtained from the Coriell Institute for Medical Research) were maintained in Eagle’s Minimum Essential Medium with Earle’s salts and non-essential amino acids supplemented with 15% FBS and 1% penicillin-streptomycin.

#### Mice

C57BL/6 mice were bred and maintained in our animal facility according to institutional guidelines and with protocols approved by the Animal Studies Committee of Washington University in St. Louis.

### METHOD DETAILS

#### Plasmids

For mammalian cell expression, human TTDN1 or DBR1 were isolated by PCR from human cDNA, cloned into pENTR-3C (Invitrogen), and subcloned into pMSCV-FLAG-HA, pHAGE-CMV-FLAG, pMSCV (no tag), or pHAGE-CMV-3XHA by Gateway recombination (Brickner *et al*., 2017; Sowa et al., 2009). TTDN1 deletions were created by PCR and cloned as above. TTDN1 point mutations and the DBR1^TTDN1-NTD^ fusion were synthesized as gBlocks (IDT) using codon sequences optimal for human cell expression, and cloned into pENTR-3C. For recombinant protein expression in bacteria, cDNAs were subcloned into pET28a-Flag or pGEX-4T1. For expressing His-GFP-DBR1 or Flag-MBP-TTDN1-mCherry and its derivatives in insect cells, cDNAs were subcloned into MacroBac 438 series vectors (Gradia et al., 2017). All constructs derived by PCR or from gBlocks were verified by Sanger sequencing.

#### Cell culture and viral transduction

293T, HeLa-S, and U2OS cells (all originally from ATCC) were cultured and maintained as previously described (Brickner *et al*., 2017). Unaffected control and NP-TTD patient fibroblast cell lines were obtained from the Coriell Institute for Medical Research and were maintained in Eagle’s Minimum Essential Medium with Earle’s salts and non-essential amino acids supplemented with 15% FBS and 1% penicillin-streptomycin (Nakabayashi *et al*., 2005). Preparation of viruses, transfection, and viral transduction were performed as described previously (Brickner et al., 2017). Knockout experiments (using lentiviral-based CRISPR/Cas9) were performed by infecting cells with the indicated lentivirus and selecting with puromycin (1 μg/ml) for 48-72 hours. For experiments with pladienolide-B, cells were treated with 250 nM of the inhibitor for 24 hours.

#### CRISPR/Cas9-mediated knockouts

The U2OS TTDN1 KO cells were created using RNP-based CRISPR/Cas9 genome editing at the Genome Engineering and iPSC Center (GEiC) at Washington University, using the gRNA sequence 5’-ACTCCCGTACCCGTCTCGAG-3’. For CRISPR/Cas9 mediated lentiviral knockout of TTDN1 and DBR1 in HeLa-S cells, gRNA sequences were cloned into pLentiCRISPR-V2 (Addgene #52961). The gRNA sequences used to generate the HeLa-S knockouts were: TTDN1, 5’-TGGCTATTATTATTACCTGG-3’; DBR1 5’-AGGCGGCAAACTTCACATGA-3’. For CRISPR/Cas9 substitution of *TTDN1* exon 1, the N-terminal aromatics mutant containing 13 alanine subsitutions for the aromatics residues (TTDN1^φ→Ala^) was cloned into an rAAV donor. The following guide RNAs were used to cleave the endogenous WT exon 1 sequence: 5’-AAATTCTGTCGCTGCATATC-3’ and 5-’ATATGCAGCGACAGAATTTT-3’. All knockout/knock-in clones were verified by deep sequencing and by Western blot analysis.

#### Purification of Flag-HA-TTDN1 and Flag-HA-DBR1 complexes and MS/MS analysis

Affinity purification of TTDN1 and DBR1 were performed as previously described, with minor modifications (Dango et al., 2011). pMSCV-Flag-HA-empty vector, TTDN1 or DBR1 retrovirus was transduced into HeLa-S cells to achieve stable expression of Flag-HA-TTDN1 or DBR1, respectively. Nuclear extract was prepared from the stable cell lines and the TTDN1 or DBR1 complexes were purified using anti-Flag resin (M2; Sigma) in TAP buffer (50 mM Tris-HCl pH 7.9, 100 mM KCl, 5 mM MgCl_2_, 10% glycerol, 0.1% NP-40, 1 mM DTT, and protease inhibitors). After elution in 1.0 mL TAP buffer plus 0.4 mg/mL Flag peptide (Sigma), the complexes were TCA precipitated, and associated proteins were identified by liquid chromatography-MS/MS at the Taplin Mass Spectrometry Facility (Harvard Medical School) using an LTQ Orbitrap Velos Pro ion-trap mass spectrometer (Thermo Fisher Scientific) and Sequest software (Sowa *et al*., 2009).

#### Immunoprecipitation and Western blotting

Immunoprecipitation of Flag- or HA-tagged GFP, TTDN1, TTDN1 mutants, or DBR1 was performed by transient transfection of constructs into 293T cells using Transit293 reagent (Mirus Bio). Cells were collected, washed in 1X PBS, and frozen at −80 °C. Pellets were resuspended in IP lysis buffer (50 mM Tris, pH 7.9, 300 mM NaCl, 10% glycerol, 1% Triton X-100, 1 mM DTT, and protease inhibitors), lysed by sonication, incubated at 4°C with rotation, and spun at 20,000 x g for 30 minutes at 4°C. An equal volume of IP lysis buffer containing no salt was added (final concentration of NaCl was 150 mM). Lysates were then incubated with anti-Flag (M2; Sigma) resin or anti-HA resin (Santa Cruz sc-7392) for 3-4 hrs at 4°C with rotation. The beads were washed extensively with IP lysis buffer containing 150 mM NaCl, and bound material was eluted with 0.4 mg/ml Flag peptide (Sigma) or with Laemmli buffer and analyzed by SDS-PAGE. For DBR1^TTDN1-NTD^ rescue experiments, cells were transduced with the indicated pHAGE-CMV lentiviral vectors. For immunoprecipitation of Flag-tagged GFP and DBR1^TTDN1-NTD^ fusion proteins from HeLa-S cells, virally transduced cells were selected with 5 μg/ml blasticidin for 48-72 hours, then collected, washed in 1X PBS, and frozen at −80 °C. Immunoprecipitation was then performed as above.

Endogenous immunoprecipitation of DBR1 was carried out from 293T cells by collecting and freezing the cells at −80 °C as above (Soll et al., 2018). Cell pellets were resuspended in TAP buffer containing 300 mM KCl, lysed by sonication, and spun at 20,000 x g for 10 minutes at 4°C. IP lysis buffer containing no salt was added to bring the final concentration of KCl to 100 mM. Samples were pre-cleared by incubation with protein A/G beads (Santa Cruz Biotechnology) with rotation at 4°C. After centrifugation, the supernatant was then incubated with equal amounts of control IgG or DBR1 antibodies at 4°C overnight with rotation. Protein A/G beads were then added and rotated at 4°C for 1 hr. The samples were then centrifuged and washed extensively with TAP buffer. Bound material was eluted with Laemmli buffer and analyzed by Western blotting.

#### Size exclusion chromatography

Nuclear extracts from control or TTDN1 KO HeLa-S cells were directly applied to a Superose 6 Increase 10/300 GL column on an AKTA Pure or AKTA go FPLC (Cytiva) equilibrated with TAP buffer. Fractions (1.0 mL each) were collected and concentrated using StrataClean Resin (Agilent). Proteins were then eluted with Laemmli buffer and analyzed by Western blotting.

#### Recombinant protein purification

For recombinant purification of GST-TTDN1, the baculovirus vector was produced using the Bac-to-Bac expression system (ThermoFisher Scientific). Amplified baculovirus was used to infect Sf9 cells and harvested after 72 hours. The cells were lysed by resuspending in Buffer L (20 mM Tris pH 7.3, 150mM NaCl, 8% glycerol, 0.2% NP-40, 0.1% TritonX-100, 2mM ϕ3-Mercaptoethanol plus protease inhibitors). Cells were lysed by sonication, then rotated at 4°C for 30 minutes. Extract was cleared by centrifugation, then added to washed Glutathione-Sepharose beads. After rotation 4°C for 2h, beads were extensively washed in Buffer L, and eluted in Buffer L plus 10mM Glutathione for 20min 4°C with rotation. Protein was dialyzed into TAP Wash buffer overnight at 4°C.

Rosetta (DE3) cells expressing His-Flag-DBR1 were resuspended in His-lysis buffer (50 mM Tris-HCl pH 7.3, 250 mM NaCl, 0.05% Triton X-100, 3 mM β−ME, 30 mM imidazole, and protease inhibitors) and lysed by sonication 3x for 30sec at 20% power. Extract was centrifuged at 12,300 x *g* for 15min 4°C, then supernatant was incubated with Nickel-NTA beads and eluted for 20 minutes at 4°C with 300 μl His-lysis buffer containing 400 mM imidazole. Protein was dialyzed into TAP Wash buffer overnight at 4°C.

Sf9 cells expressing Flag-MBP-TTDN1-mCherry and Flag-MBP-TTDN1 Aro>Ala-mCherry were harvested and frozen at −80°C. Pellets were resuspended in MBP Lysis buffer (50mM Tris-HCl pH 7.9, 500mM NaCl, 5% glycerol, 0.5mM DTT, 1mM PMSF, and protease inhibitors). After douncing, the cell extracts were further lysed by sonication on ice at 25% amplitude for 3 minutes (30 seconds on, 30 seconds off) and centrifuged at 12,000 *x g* for 30 minutes at 4°C. The supernatant was incubated with Hi-Flow amylose resin (NEB) for 1hour at 4°C, then washed extensively in MBP Lysis Buffer. Elution using MBP lysis buffer was performed in the presence of 12.5 μl Precission protease (ThermoFisher, 2U/μl) at room temperature for 1 hour with rotation. Washed glutathione-Sepharose resin (Sigma) were added for 15 minutes to remove remaining Precission protease. For *in vitro* droplet assays performed in the absence of PEG, TTDN1-mCHerry eluates were concentrated using Amicon Ultra-15 Centrifugal Filters (Millipore). Sf9 cells expressing His-GFP-prp-DBR1 were harvested and frozen at −80°C. Pellets were resuspended in 30 mL Buffer L (50 mM Tris pH 7.3, 500 mM NaCl, 8% glycerol, 0.2% NP-40, 0.1% Triton X-100, 25 mM Imidazole, 1 mM β-Mercaptoethanol). An additional 30 mL Buffer L was added prior to 30 minutes of rotation at 4°C to complete cell lysis. Extract was centrifuged at 12,300 x *g* for 10 minutes, then supernatant was incubated with Ni-NTA beads and eluted with Buffer L containing 400 mM imidazole. After dialysis into TAP buffer, protein was concentrated using Amicon Ultra-15 Centrifugal Filters (Millipore), and then sample was directly applied to a Superose 6 Increase 10/300 GL column on an AKTA Pure FPLC (Cytiva) equilibrated with TAP buffer. 1 mL fractions were collected and analyzed by Coomassie Blue staining. Peak fractions were kept and stored at −80°C.

#### *In vitro* condensate formation assays

Recombinant mCherry (Biovision; #4993), TTDN1-mCherry, TTDN1 Aro>Ala-mCherry, and/or GFP-DBR1 were rapidly thawed at 37°C, then diluted in buffer containing 50 mM Tris-HCl pH 7.9, 5% glycerol, and the indicated NaCl and PEG-8000 concentrations. Samples were mixed by brief vortexing, incubated at room temperature for 10 minutes, and visualized using an Olympus fluorescence microscope (BX-53) using an UPlanS-Apo 100×/1.4 numerical aperture oil immersion lens and cellSens Dimension software. Live imaging for droplet fusion and fission were visualized with Olympus fluorescence microscope (IX-83) using an UPlanS-Apo 100×/1.4 numerical aperture oil immersion lens and cellSens Dimension software.

#### Phase separation simulations

All-atom simulations were run using the CAMPARI simulations engine (http://campari.sourceforge.net/) and ABSINTH implicit solvent model (abs_opls_v3.2) (Vitalis and Pappu, 2009). Simulation setup was identical to that performed previously for the 135-residue low-complexity domain from hnRNPA1 (Martin *et al*., 2020). Thirty independent simulations were run for 150 M steps each with conformations saved every 100 K steps, generating a final ensemble of 42,000 conformations. The large interval of 100 K steps between conformation output is necessary given the large size of the IDR and ensures that local conformational correlations are minimized. All-atom simulations were analyzed using SOURSOP (https://soursop.readthedocs.io/). Contacts were defined as two residues with heavy atoms within 5.0 Angstroms of one another. Secondary structure was calculated based on DSSP. Coarse-grained simulations were run using PIMMS (https://zenodo.org/record/3588456) with identical parameters applied previously for hnRNPA1 simulations (Martin *et al*., 2020). Binodals were calculated from simulation analysis using the radial droplet density to measure the high-concentration arm of the binodal. Sequence analysis (including patterning analysis) was performed using localCIDER (Holehouse et al., 2017).

#### Protein binding assays

All *in vitro* GST-protein binding assays were performed as described previously with minor modifications (Mosammaparast et al., 2013). Briefly, 6 μg of the indicated GST-tagged protein was incubated with 30 μl of blocked glutathione-Sepharose beads and 2 μg of His_6_-Flag-DBR1 in TAP buffer containing 1% BSA in a total volume of 100 μl. After incubation at 4°C with rotation for 1 hour, beads were washed extensively using TAP buffer, followed by a final wash in 1X PBS. Bound material was eluted using Laemmli buffer and analyzed by SDS-PAGE and Western blotting. For mCherry-protein binding assays, anti-mCherry Nanobody Affinity Gel (Biolegend) were blocked with BSA overnight at 4°C in 1X TAP buffer and 3.3% BSA. 10 μg of the indicated mCherry protein was incubated with 30 μl blocked Anti-mCherry Nanobody affinity gel in TAP buffer in a total volume of 100 μl. After incubation at 4°C with rotation for 1 hour, beads were washed extensively using TAP buffer. After addition of 1μg of GFP-DBR1, samples were incubated at 4°C with rotation for 1 hour, washed extensively using TAP buffer followed by a final wash in 1X PBS. Samples were eluted using Laemmli buffer, analyzed by SDS-PAGE and Western blotting or Coomassie Blue staining.

#### RNA-Seq and data analysis

RNA was purified from cell lines using the Qiagen miRNeasy mini kit to accomodate small RNA isolation (#217004). Samples for small RNA-sequencing were prepped with TruSeq Small RNA library preparations kits; otherwise all other samples were prepared according to library kit manufacturer’s protocol, indexed, pooled, and sequenced on an Illumina NovaSeq 6000 2×150bp with the Genome Technology Access Center at Washington University in St. Louis, typically yielding 200 million paired-end reads per sample.

Total RNA isolation from mouse cortex was carried out as previously described (Christian et al., 2020). In brief, cerebral cortex was dissected in ice-cold PBS from 5 female *Ttdn1^Δ/Δ^* and 5 WT littermates at 8 weeks of age. RNA was purified from cortex using the Qiagen miRNeasy mini kit. Samples were prepared according to library kit manufacturer’s protocol, indexed, pooled, and sequenced on an Illumina NovaSeq 6000 2×150bp with the Genome Technology Access Center at Washington University in St. Louis, typically yielding 200 million paired-end reads per sample.

Basecalls and de-multiplexing were performed with Illumina’s bcl2fastq software and a custom Python demultiplexing program with a maximum of one mismatch in the indexing read. RNA-Seq reads were then aligned to Ensembl GRCh38.76 or Ensembl GRCm38.76 assembly for human or mouse samples, respectively, with STAR version 2.7.9a (Dobin et al., 2013). Gene counts were derived from the number of uniquely aligned unambiguous reads by Subread:featureCount version 2.0.3 (Liao et al., 2014). Isoform expression of known Ensembl transcripts were quantified with Salmon version 1.5.2 (Patro et al., 2017). Sequencing performance was assessed for the total number of aligned reads, total number of uniquely aligned reads, and features detected. The ribosomal fraction, known junction saturation, and read distribution over known gene models were quantified with RSeQC version 4.0 (Wang et al., 2012).

All gene counts were then imported into the R/Bioconductor package EdgeR (Robinson et al., 2010) and TMM normalization size factors were calculated to adjust for samples for differences in library size. Ribosomal genes and genes not expressed in the smallest group size minus one sample greater than one count-per-million were excluded from further analysis. The TMM size factors and the matrix of counts were then imported into the R/Bioconductor package Limma (Ritchie et al., 2015). Weighted likelihoods based on the observed mean-variance relationship of every gene and sample were then calculated for all samples and the count matrix was transformed to moderated log 2 counts-per-million with Limma’s voomWithQualityWeights (Liu et al., 2015; Luo et al., 2009). The performance of all genes was assessed with plots of the residual standard deviation of every gene to their average log-count with a robustly fitted trend line of the residuals. Differential expression analysis was then performed to analyze for differences between conditions and the results were filtered for only those genes with Benjamini-Hochberg false-discovery rate adjusted p-values less than or equal to 0.05. For each contrast extracted with Limma, global perturbations in known Gene Ontology (GO) terms, MSigDb, and KEGG pathways were detected using the R/Bioconductor package GAGE (Luo *et al*., 2009) to test for changes in expression of the reported log 2 fold-changes reported by Limma in each term versus the background log 2 fold-changes of all genes found outside the respective term.

#### qRT-qPCR

RNA was extracted using the QIAGEN miRNeasy Mini Kit (#217004). Reverse transcription was performed on 2μg purified RNA using the High-capacity cDNA reverse transcription kit with RNase inhibitor (ThermoFisher) using poly(dT) primers. SYBR Green JumpStart Taq Ready Mix (Sigma S9194) with used with qPCR using the QuantStudio 6 Flex Real-Time PCR System (Applied Biosystems). Relative quantification was performed using the 2^-1111Ct^ method.

#### rMATS

rMATS turbo v4.1.1 was used to detect the splicing events and significant splicing differences between TTDN1 or DBR1 knockout and control samples, including patient fibroblast samples and mouse samples. “Positive” and “negative” indicates inclusion and exclusion of splicing event relative to control transcript. For an event to be considered for any downstream analysis we required that each isoform was supported by at least 5 reads in half of the samples. Differentially spliced events were required to have an absolute difference in inclusion level greater than 10% and a false discovery rate less than 10% (Shen et al., 2014) (https://github.com/Xinglab/rmats-turbo).

#### Gene length analysis

Lengths of transcripts (CDSs plus UTRs) and numbers of exons for the human genome assembly were obtained by intersecting the information from file knownGene.txt (https://hgdownload.soe.ucsc.edu/goldenPath/hg38/database/, June 2020) with a custom BioMart Ensembl file of GRCh38 (May 2020) using in-house scripts. One-exon transcripts were excluded. The log10 of both transcript length and exon number were used for the analyses. RNA-Seq data were taken from the list of differentially expressed transcripts displaying significant Benjamini-Hochberg false-discovery rates, using various log2 fold-changes cutoffs as specified, and comparing with the remaining transcripts. For the line plots of transcript length or exon number versus log2 fold-changes we binned the ranked data with stride = 200 (i.e. 200 transcripts per bin) and displacement = 40 (overlap) unless otherwise specified. For each data point, error range corresponds to the standard error. For the mouse genome assembly, we used Ensembl GRCm38.76. We obtained transcript lengths and exon numbers using Daren Card’s utility “genestats.sh” at https://gist.github.com/darencard/fcb32168c243b92734e85c5f8b59a1c3. The “bgzip” and “tabix” utilities were downloaded from http://www.htslib.org/download/ (version htslib-1.12); “bedtools” was version 2.29.2. The R libraries “ggplot2”, “ggpubr”, and “grid” along with the Wilcoxon test were used to construct box plots. These show the median (unless specified otherwise), interquartile range (IQR) from Q1 (25^th^ percentile) to Q3 (75^th^ percentile), whiskers extending from Q1 – 1.5xIQR (minimum) to Q3 + 1.5xIQR (maximum), and outliers as dots.

#### Lariat identification by splice site matching

A lariat mapping pipeline was developed based on the method described in Pineda and Bradley, 2018. First, reads are filtered out if they contain >5% ambiguous characters. Then, reads are mapped to the genome, and aligned reads are discarded. A mapping index is then created based on the unaligned reads, and a FASTA file containing the first 20nt of each annotated intron in the transcriptome is mapped to the unaligned reads. Reads are then identified where only one 5’ splice site maps to them and the alignment has no mismatches or indels. These reads are then trimmed of the sequence from the start of the 5’ SS alignment to the end of the read, and reads were the trimmed portion is <20nt are filtered out. The remaining trimmed reads are mapped to an index built from the last 250nt of every annotated intron. The trimmed read alignments are then filtered to only consider those with <=5 mismatches, <=10% mismatch rate, and no more than one indel of <=3nt. Then, for each trimmed read the highest scoring alignment was chosen after restricting to alignments in the same gene as the 5’ SS alignment and those with the expected inverted mapping order of the 5’ and 3’ segments. The end of this highest scoring alignment is then taken to be the branchpoint of the lariat the read is derived from.

#### Immunofluorescent microscopy

U2OS cells expressing pHAGE-CMV-3XHA-TTDN1 were washed with 1X PBS before fixation with 3.2% paraformaldehyde. The cells were then washed with immunofluorescence wash buffer (1× PBS, 0.5% NP-40, and 0.02% NaN_3_), then blocked with immunofluorescence blocking buffer (immunofluorescence wash buffer plus 10% FBS) for 30 minutes. HA (Santa Cruz sc-805) antibody was diluted in immunofluorescence blocking buffer at 1:300 overnight at 4°C. After staining with Alexa Fluor 488 secondary antibody (Millipore) and Hoechst 33342 (Sigma-Aldrich), samples were mounted using Prolong Gold mounting medium (Invitrogen). Epifluorescence microscopy was performed on an Olympus fluorescence microscope (BX-53) using an UPlanS-Apo 100×/1.4 numerical aperture oil immersion lens and cellSens Dimension software. Raw images were exported into Adobe Photoshop, and for any adjustments in image contrast or brightness, the levels function was applied.

#### Targeting mouse *Ttdn1* locus and animal husbandry

*Ttdn1^Δ/Δ^* and *Ttdn1^M143V/M143V^* mice were created using CRISPR/Cas9 technology at the Washington University Genome Engineering and iPSC Center as previously described (Zhao et al., 2018). Female C57BL/6 mice were super-ovulated using 5 IU of Pregnant Mares Serum Gonadotropin followed by 5 IU of Human Chorionic Gonadotropin 48 hours later. The females were then mated to C57BL/6 male mice and day 0.5 embryos were isolated the morning after mating. The fertile single cell embryos underwent pronuclear micro-injection delivering the CRISPR gRNA (5’-TTCAATGCTTGAAGACCCTTNGG-3’) mixed with RNA encoding Cas9 and donor DNA. The concentration of the injection mix was 5 ng/μl gRNA with 10 ng/μl Cas9 RNA. Tail samples were taken from pups and deep sequencing was performed to identify animals carrying indels, and the exact modification that occurred. The founder male mouse with the *Δ34* allele, or the *M143V* knockin allele, was selected and mated to a C57BL/6 female to isolate the allele, and heterozygous progeny were mated to generate the homozygous mutant mice.

All animal protocols were approved by the Institutional Animal Care and Use Committee and the Animal Studies Committee of Washington University in St. Louis, and in accordance with guidelines from the National Institutes of Health (NIH). Mice were housed in a room on a 12:12 hour light/dark cycle, with controlled room temperature (20-22°C) and relative humidity (50%). Home cages measured 28.5 cm x 17.5 cm x 12 cm and were supplied with corncob bedding and standard laboratory chow and water. All mice were group-housed and adequate measures were taken to minimize animal pain or discomfort.

#### Behavioral analysis

The behavioral procedures were similar to previously described methods (Maloney et al., 2019), except for the marble burying test, for which the general procedures described in (Cheng et al., 2018) were used. All behavioral testing was conducted during the light cycle, by an experimenter “blinded” to experimental group status of each mouse. The order of the tests was the same as described below, which reflects attempts to minimize “carry-over” effects across measures by conducting the most stressfμl measures last in the series. Only one test was conducted per day. All equipment was cleaned with 2% chlorhexidine diacetate or 70% ethanol between animals. Behavioral studies were conducted in two different cohorts of mice: cohort 1 – *Ttdn1^+/+^* =10 (4M; 6F); *Ttdn1^Δ/Δ^* =10 (4M; 6F); and cohort 2 - *Ttdn1^+/+^* =8 (4M; 4F); *Ttdn1^Δ/Δ^* =9 (4M; 5F). Results presented are from analyzing the combined data set from the two different cohorts of mice. Analysis of the combined data set was carried out to increase statistical power for evaluating main effects of Genotype and Sex and their interactions. Behavioral testing involved using identical protocols and test sequences for each cohort. Moreover, we conducted “analysis screens” on all the behavioral data using an ANOVA model containing Genotype and Cohort as between-subjects variables. These models were used to determine whether any Genotype x Cohort interactions were present in order to judge the appropriateness of combining the data sets. These analyses did not produce any significant Genotype x Cohort interactions per se. However, in two instances, stationary rod (rotarod) and place path length (Morris water maze), 3-way interactions including a Genotype x Cohort along with the main dependent variable were observed, but this was of no consequence since there were no differences between groups with regard to these two variables. Note that there was one difference between the testing of the two cohorts. Specifically, only the *Ttdn1^Δ/Δ^* and Control mice from cohort 2 were evaluated on acoustic startle and prepulse inhibition of startle (PPI). This test was conducted one week after the mice were assessed on conditioned fear to determine the likelihood that the significant impairment on the auditory cue test exhibited by the TTDN1 mice was due to severe auditory deficits

#### Rotarod

The *Ttdn1^Δ/Δ^* and WT mice were tested on the rotarod (Economex, Columbus Instruments, Columbus, OH) to assess fine motor control. Each mouse was placed on a 9.5 cm section of grooved rod measuring 3.81 cm in diameter surrounded by plastic walls and elevated 52 cm from the floor. Mice received five trials on three test days, where each test session was separated by 4 days to minimize motor learning. The rod was stationary for trial 1 and continuously rotating at 2.5 rpm for trials 2 and 3 for 60 s. The rod accelerated by 0.13 rpm per second for trials 4 and 5 for 180 s. Time the mouse remained on the rod served as the dependent variable. Note that the one-hour locomotor activity test, sensorimotor battery, and marble burying test were conducted during days intervening between rotarod test sessions, although only one test was scheduled for a given day.

#### One-hour locomotor activity and sensorimotor battery

Mice were placed in transparent polystyrene enclosures (47.6×25.4×20.6cm) and movements were monitored using computerized photobeam instrumentation. General activity variables (total ambulations, number of vertical rearings) were collected along with emotionality indices (time spent, distance traveled, and entries made in a 33 x 11 cm central zone, as well as distance traveled within a 5.5 cm contiguous area around the periphery. The following day, mice were run on a battery of sensorimotor tests (walking initiation, ledge platform; pole, 60° and 90° inclined screens, inverted screens) to assess movement initiation, balance, strength, and coordination. For walking initiation, mice were placed in the center of a 21 × 21 cm square marked with tape, and the amount of time mice took to leave the square was recorded. In the ledge and platform tests, mice were placed on an elevated Plexiglas ledge (0.75 cm wide) or small circular wooden platform (3.0 cm diameter) elevated to 30 or 47 cm, respectively, and the amount of time they could remain on either apparatus was recorded. In the pole test, mice were oriented head-up with forepaws on top of a textured rod (8 mm x 55 cm) and the amount of time the mouse took to turn around and climb down the pole was measured. If the mouse fell off the pole, it was assigned a maximum score of 120 s. Inclined screen tests were performed by placing mice head-oriented down on a wire mesh grid (16 x 10 cm) elevated 47 cm and inclined at 60° or 90°. The time taken for the mouse to turn 180° and climb to the top of the wire mesh was then measured. Inverted screen tests began identically to inclined screen tests described above, except that the screen was inverted 180° after ensuring the mouse had a secure grip. The amount of time the mouse could remain on the screen was recorded. Each test lasted a maximum of 60 s, except for the pole measure (120 s). Means from two trials per test per mouse were used in all analyses.

#### Marble burying

Species-specific, compulsive digging behavior was evaluated in the mice using the marble burying test employing a procedure generally similar to previously-described methods (Maloney et al., 2018). A rat cage was filled with aspen bedding to a depth of 3 cm served as the apparatus. Twenty marbles were placed on top of the bedding in a 5 × 4 evenly spaced configuration. The test began by placing a mouse in the center of the chamber and allowing it to freely explore and dig for 30 min under normal laboratory lighting conditions. An acrylic lid containing air holes was placed on top of the cage to prevent mice from escaping. After 30 min, the mouse was returned to its home cage. Two observers counted the number of marbles not buried (less than two-thirds of the marble was covered with bedding). The number of marbles buried was then determined, and the average of the two scores was used in the analysis. After the marbles were counted, the bedding was disposed of and the cage and marbles were cleaned with 2% chlorhexidine diacetate.

#### Morris Water Maze (MWM)

Spatial learning and memory were assessed using the MWM three days after completion of the sensorimotor battery. A computerized tracking system (ANY-maze; Stoelting) recorded the swim path of the mouse to the escape platform and quantified path length and latency to the escape platform, and swimming speeds during cued, place, and probe trials conducted in a pool of opaque water. Mice were first tested on the cued condition to assess whether they had any demonstrable nonassociative deficits (e.g., visual or sensorimotor disturbances) that might affect subsequent performance during the place (spatial learning) trials. For the cued trials, the escape platform was submerged beneath the surface of the water, but its location was denoted by a red tennis ball atop a rod, which was attached to the escape platform and served as a visual cue. Cued trials were conducted four times per day for two consecutive days, for a total of eight trials with an inter-trial interval (ITI) of 30 min and a 60 s maximum per trial. To limit spatial learning during cued trials, the location of the platforms was varied across trials in the presence of very few distal spatial cues. Cued trial performance was analyzed as four blocks of two trials each. Three days after completing the cued trials, spatial learning was assessed during the place condition. For the place trials, the platform was hidden beneath the surface of the opaque water and its location was kept constant across all trials in the target quadrant, with several salient distal cues being present. Acquisition training involved releasing a mouse from each of the pool quadrants for each trial, with the sequence of quadrants being pseudorandomly determined for each test session. Two blocks of 2 consecutive trials each were performed over five days, with 60 s maximum per trial and an ITI of 30 s, during which time a mouse was allowed to remain on the platform. Blocks were separated by approximately 2 h. The place trials data were analyzed over five blocks of four trials, each block representing one day of training. A 60 s probe trial was administered about 1 h after the last place trial on the fifth day when the platform was removed, and the mouse was released into the maze from the quadrant opposite where the platform had been located. The amount of time the mouse spent searching in each quadrant of the pool, as well as the number of times it crossed over the exact location where the platform had been located (platform crossings) were recorded.

#### Conditioned fear

Three days after completing testing in the MWM, the mice were assessed on Pavlovian fear conditioning. The procedure involved placing a mouse into a Plexiglas conditioning chamber (26cmx18cm x18cm; Med-Associates), that contained distinct visual, tactile, and olfactory cues. Freezing behavior was assessed for a 2-min baseline period prior to tone-shock training. Three minutes after being placed in the conditioning chamber, and every 60 s thereafter, mice were exposed to 3, tone-shock pairings. Each pairing consisted of 20 s of broadband white noise presented at 80 dB (conditioned stimulus; CS), with a 1.0 mA continuous foot shock (unconditioned stimulus; UCS) presented during the last second of the tone. Mice were placed back into the same conditioning chamber the following day and freezing behavior was measured over an 8-min period. One day later, mice were placed into a chamber that contained a different set of cues. Freezing behavior was recorded for a 2-min altered context baseline period, after which mice were assessed on the auditory cue test, which involved the presentation of the tone (CS) over an 8 min period. Freezing behavior was quantified using FreezeFrame image analysis software (Actimetrics), where freezing was defined as no movement beyond that associated with breathing. Data are presented as a percentage of time spent freezing, relative to the total duration of the trial. Shock sensitivity was evaluated after fear conditioning using previously described procedures.

#### Acoustic startle/prepulse inhibition of startle (PPI)

After the completion of testing the second cohort of mice, the decision was made to evaluate the second cohort on the acoustic startle response and prepulse inhibition of the startle response (PPI) to provide some information on the possibility that deafness or severe auditory deficits were responsible for the impaired auditory cue performance during conditioned fear testing (day 3) of the *Ttdn1^Δ/Δ^* mice. Thus, one week after completing the conditioned fear test for cohort 2, sensorimotor reactivity and gating and the general intactness of the auditory system were evaluated in the mice by quantifying the magnitude of their acoustic startle response and PPI (Hamilton Kinder, LLC), using methods similar to those previously described (Cheng, 2018). Specifically, responses to a 120 dB auditory stimulus pulse (40 ms broadband burst) and PPI (response to a prepulse plus the startle pulse) were measured concurrently in the mice using Kinder Scientific Startle Reflex chambers (Poway, CA, USA). A total of 20 startle trials were presented over a 20 min test period during which the first 5 min served as an acclimation period when no stimuli above the 65 dB white noise background were presented. The session began and ended by presenting 5 consecutive startle (120 db pulse alone) trials unaccompanied by other trial types. The middle 10 startle trials are interspersed with PPI trials (consisting of an additional 30 presentations of 120 dB startle stimuli preceded by pre-pulse stimuli of either 4, 12, or 16 dB above background (10 trials for each PPI trial type). Following pseudorandom presentation of all PPI and startle trials, responses to 40 ms broadband bursts at 80, 90, 100,110 and 120 dB were measured to screen for differences in auditory thresholds. A %PPI score for each trial was calculated using the following equation: %PPI= 100*(ASRstartle pulse alone - ASRprepulse+startle pulse)/ASRstartle pulse alone.

#### Neurohistology

At around 4.5 months of age, animals were perfused using 4% paraformaldehyde and brains sectioned on a vibratome at 75 microns. Sections were then stained with hematoxylin by immersing in two five-minute exchanges of 100% ethanol, 2 minutes in 95% ethanol, 2 minutes in 70% ethanol, five dips in deionized water, and eight minutes in hematoxylin (Gill’s hematoxylin stock solution, Sigma-Aldrich, GHS132-1L). This was followed by two exchanges of five dips in deionized water, ten seconds in 0.2% ammonia water, and a final five dips in deionized water. Mounted sections were then coverslipped using an aqueous mounting medium (Permanent Aqueous Mounting Medium, Bio-Rad, BUF058A). Regional volumes were then quantified using Stereo Investigator Software (Version 2020.2.3, MBF Bioscience, Williston, VT) running on a Dell Precision Tower 5810 computer connected to a QImaging 2000R camera and a Labophot-2 Nikon microscope with electronically driven motorized stage.

---

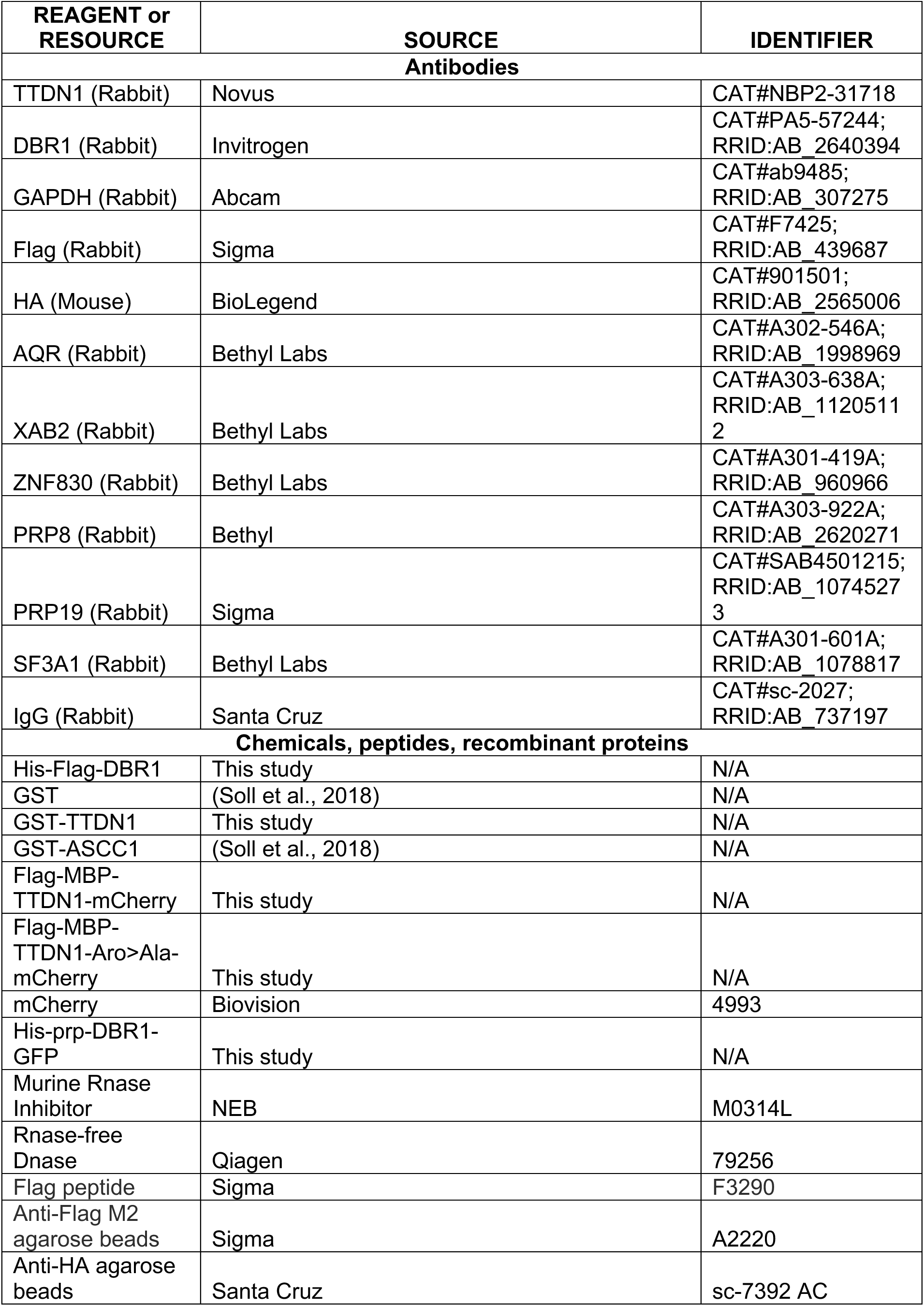
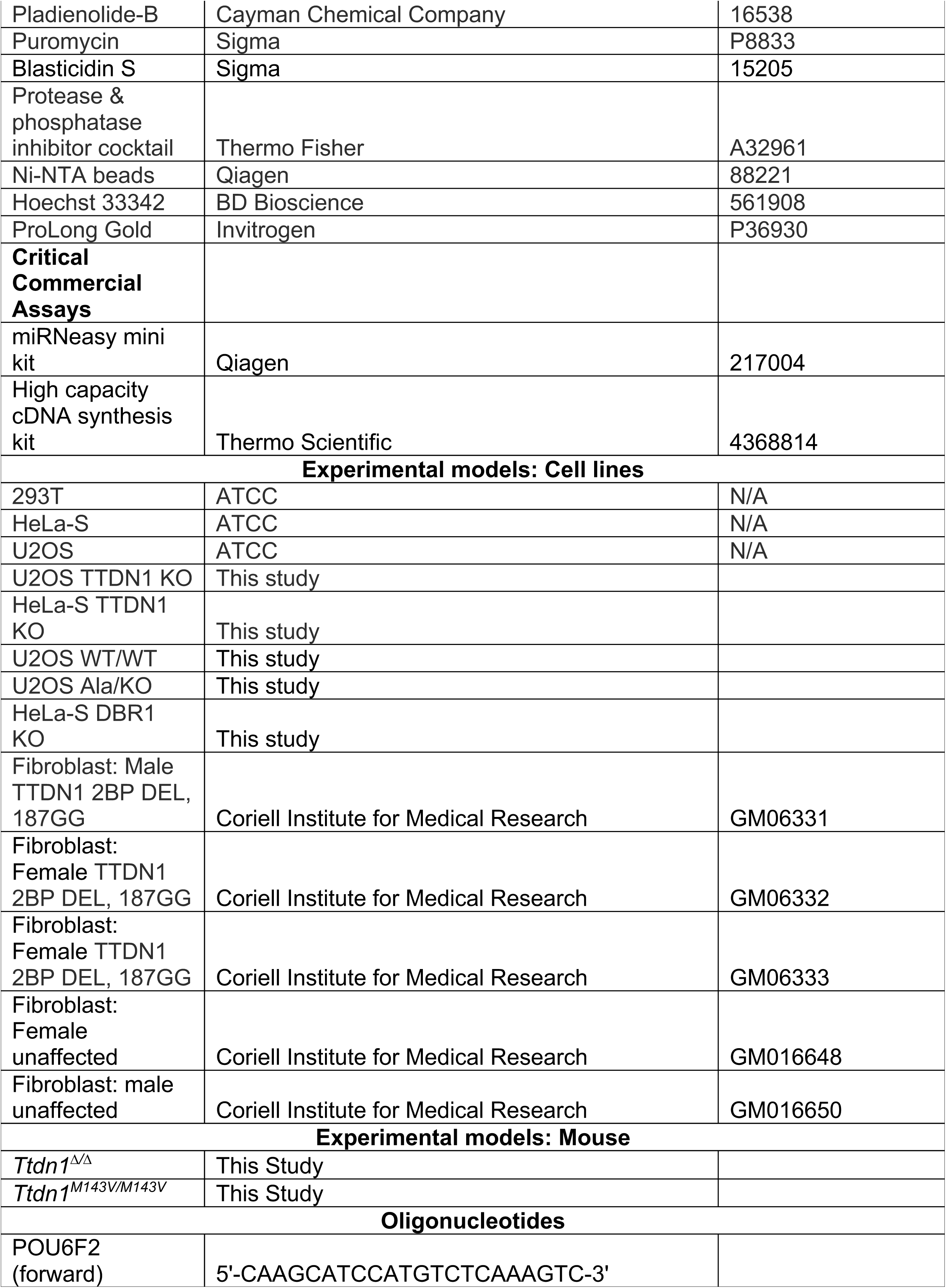
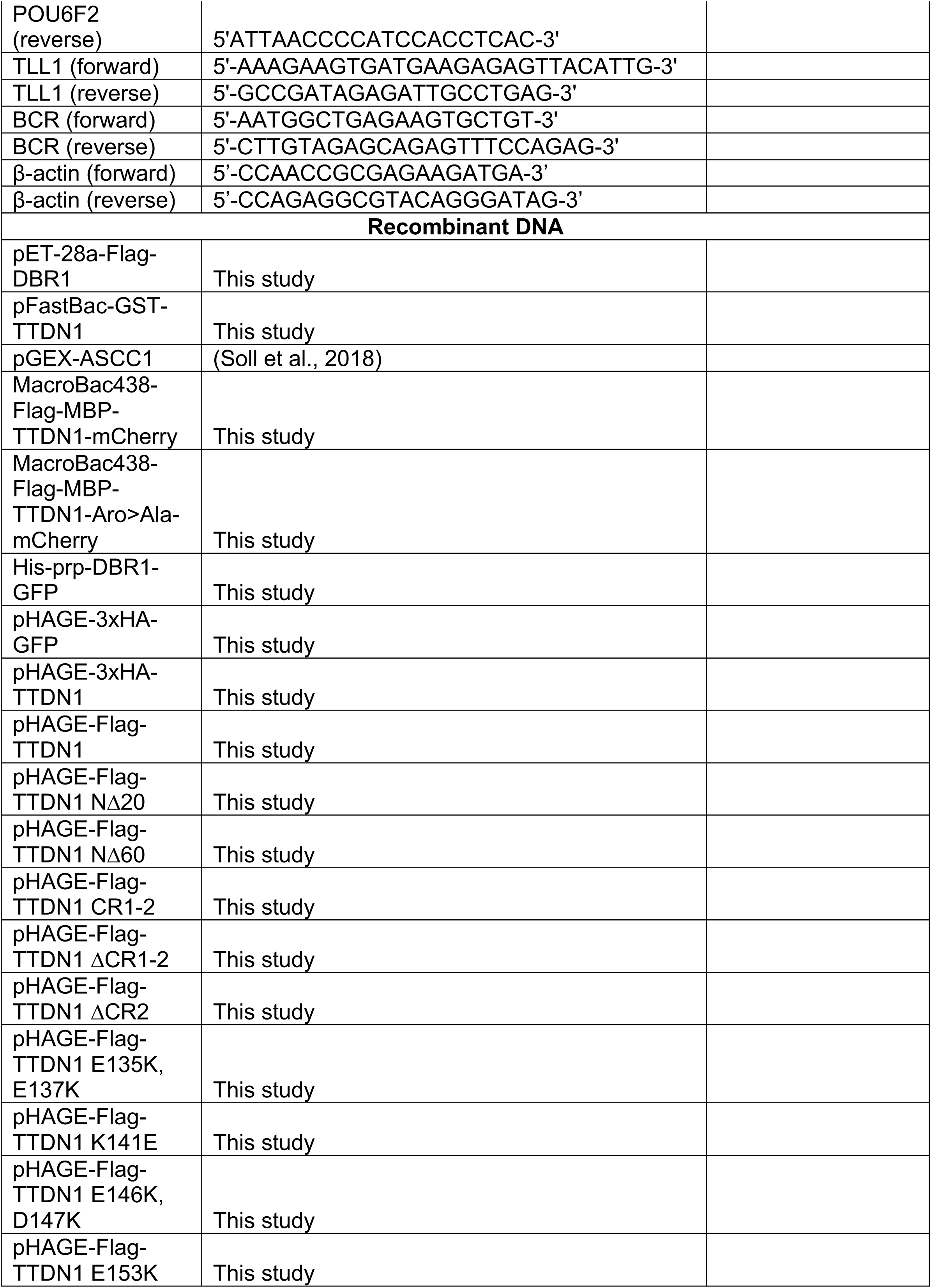
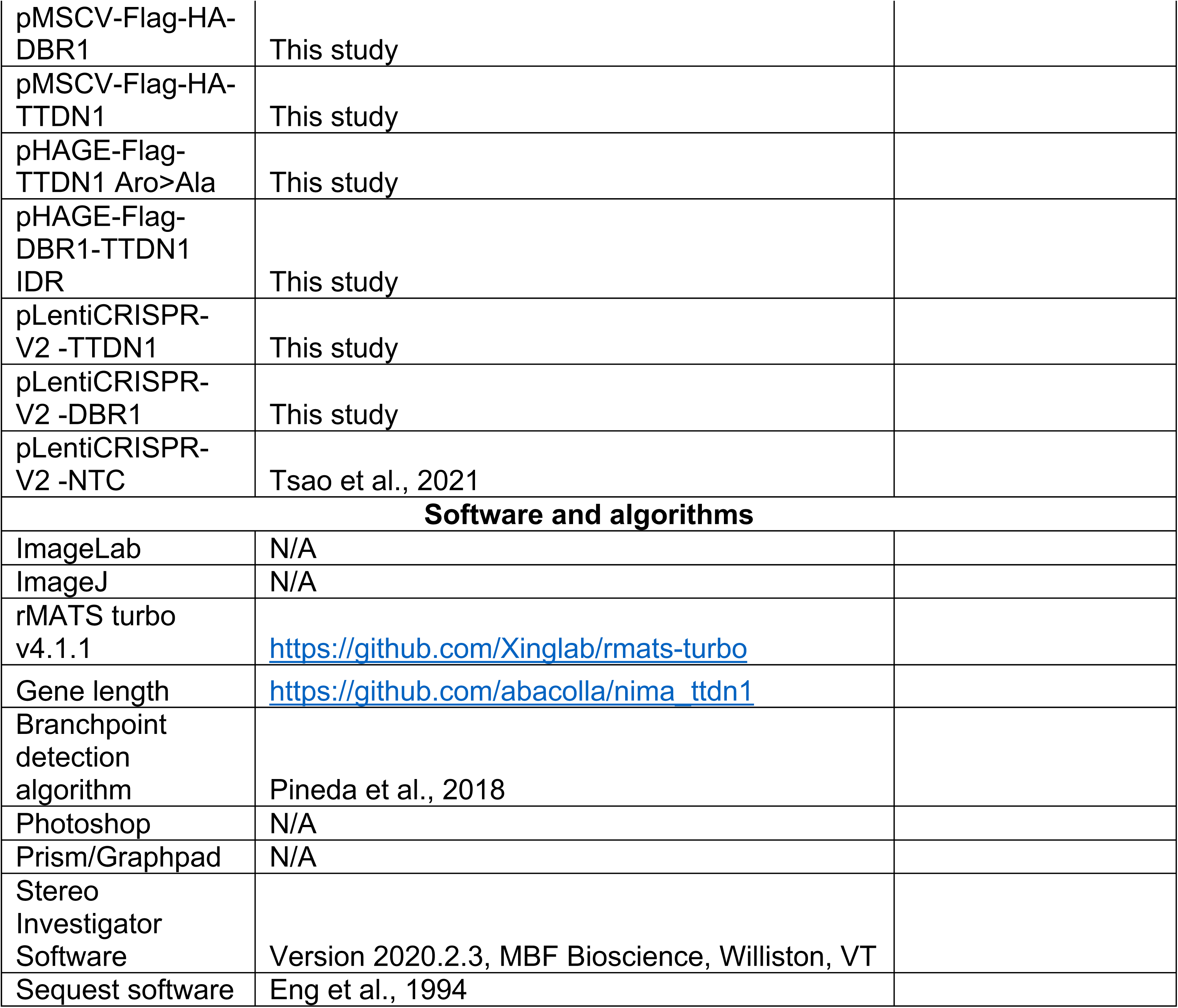

## Supplemental Figure Legends

**Supplemental Figure S1.**
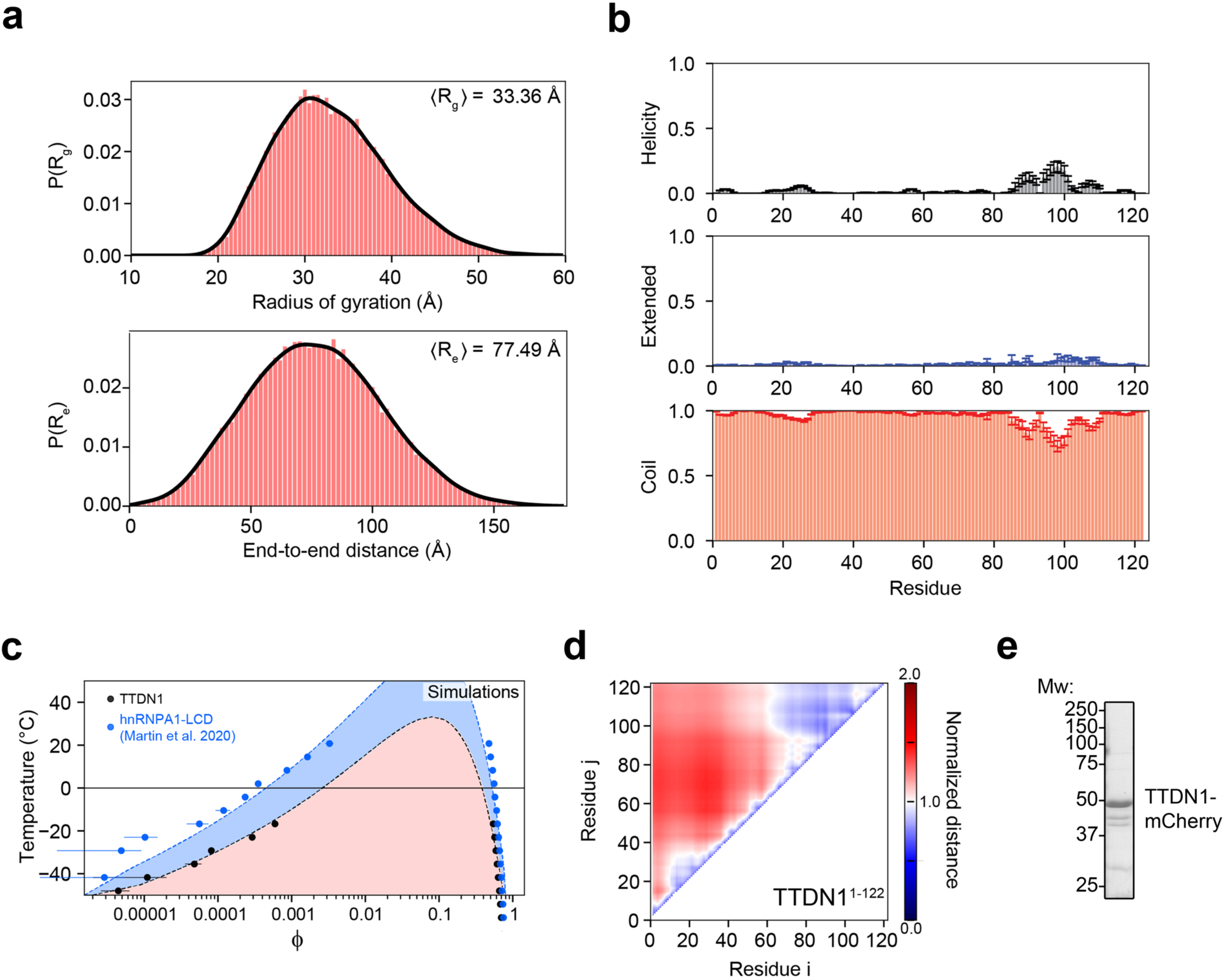
TTDN1 as a putative phase-separating molecule (related to Figure 1). **(a)** The radius of gyration (*R_g_*) and end-to-end distance (*R_e_*) offer polymeric metrics used to quantify the global dimensions of disordered proteins. (top) All-atom simulations reveal an expected *R_g_* for TTDN1^1-122^ of around 33 Å, suggesting the IDR is relatively expanded when compared to the *R_g_* measured for other disordered proteins of a similar length (Lazar et al., 2021). The broad smooth distribution is indicative of effective and well-resolved conformational sampling obtained by the simulations. (bottom) The end-to-end distance distribution is similarly broad and follows the expected distribution for an expanded and flexible polymer. **(b)** Secondary structure analyzed by DSSP confirms the disordered nature of TTDN1^1-122^ with the possible exception of transient residual helicity between residues 90 and 110. As per standard DSSP classification, secondary structure is defined as either (top) helicity, (middle) ‘extended’ (which includes beta-sheet and beta-strand conformations) and (bottom) coil. Quantification reflects the fraction of all simulations for which each residue is found in a given structural state. Error bars reflect standard error of the mean over thirty independent replicas. **(c)** TTDN1^1-122^ phase diagram (red) compared to that reported previously by Martin *et al*. (blue) (Martin et al., 2020). Coarse-grained simulations were run as a function of temperature and the resulting phase boundaries (binodals) show good agreement with a hypothetic phase diagram generated using Flory-Huggins theory. Error bars are the standard error of the mean as calculated over three independent simulations. **(d)** Intramolecular scaling map quantifying average inter-residue distances as normalized by the distances expected for a Gaussian chain, as described previously (Holehouse et al., 2015). **(e)** TTDN1-mCherry was purified from Sf9 cells and analyzed by SDS-PAGE, followed by Coomassie blue staining. Positions of molecular weight markers are shown on the left (Mw).

**Supplemental Figure S2.**
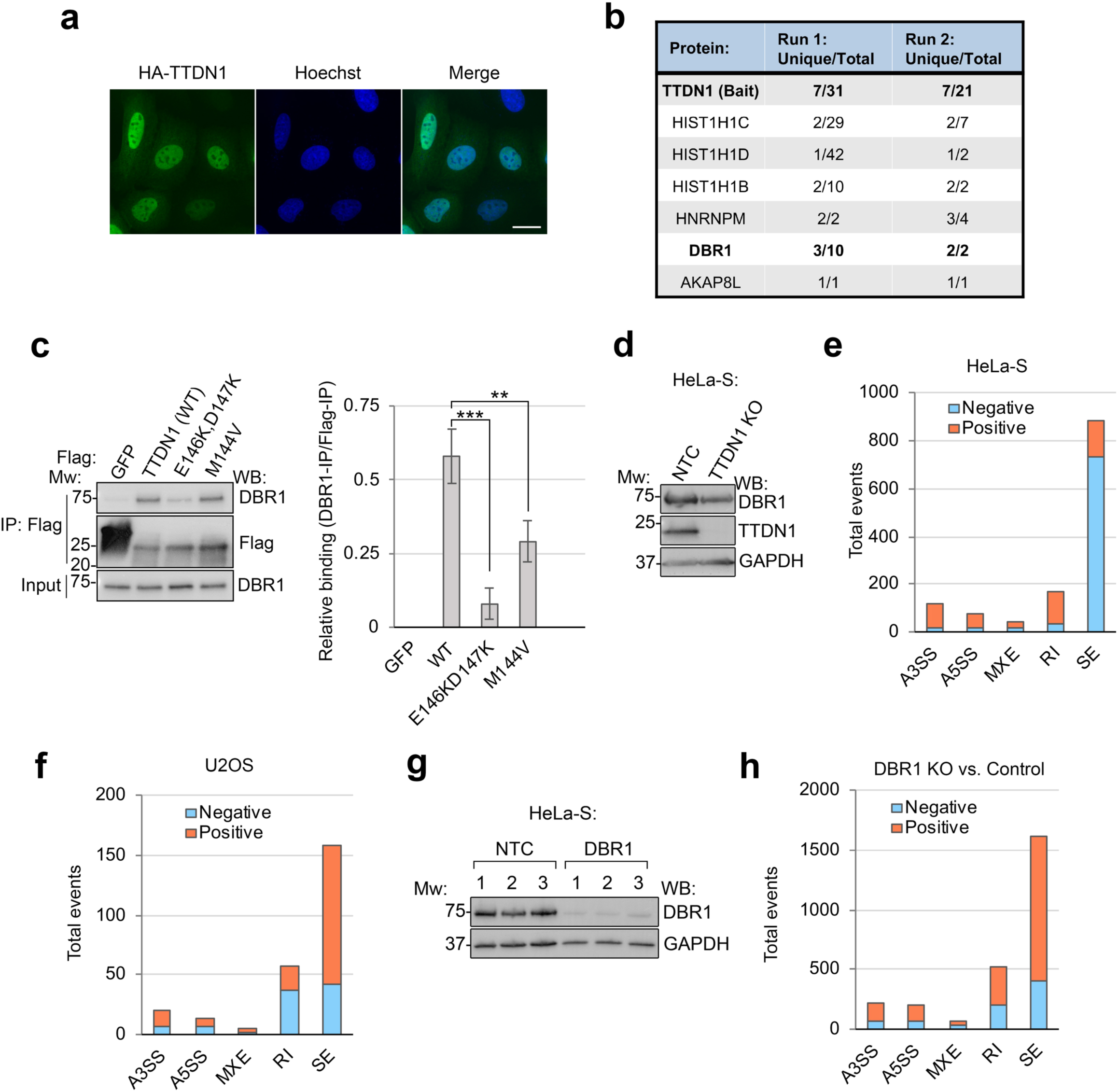
Proteomic and functional analysis of TTDN1 (related to Figure 2). **(a)** Immunofluorescence of HA-TTDN1-expressing cells. Scale bar, 20 µm. **(b)** Flag-HA-TTDN1 was isolated from HeLa-S nuclear extract using Flag immunoprecipitation. Peptides unique to the TTDN1 purification from two independent experiments were identified by LC-MS/MS. Peptide numbers for each protein are shown. **(c)** Flag IP was performed from 293T cells expressing the indicated proteins. IP and input material were analyzed by Western blot as shown. Figure representative of four independent experiments. Quantification on right. ** p<0.01, *** *p* < 0.001 by unpaired t-test. **(d)** DBR1 expression was evaluated in HeLa-S cells from **Figure 2g** by Western blot. **(e)** and **(f)** Alternative 3′ and 5′ splice sites (A3SS, A5SS), mutually exclusive exons (MXE), retained introns (RI), and skipped exons (SE) were quantified using rMATS analysis on n=3 technical replicate RNA-Seq samples from **Figure 2h-i**. **(g)** DBR1 was targeted using CRISPR/Cas9 in HeLa-S cells. Three separate knockout pools were analyzed by Western blot using antibodies as shown. **(h)** Splicing event analysis was performed using rMATS from the DBR1 KO cells.

**Supplemental Figure S3.**
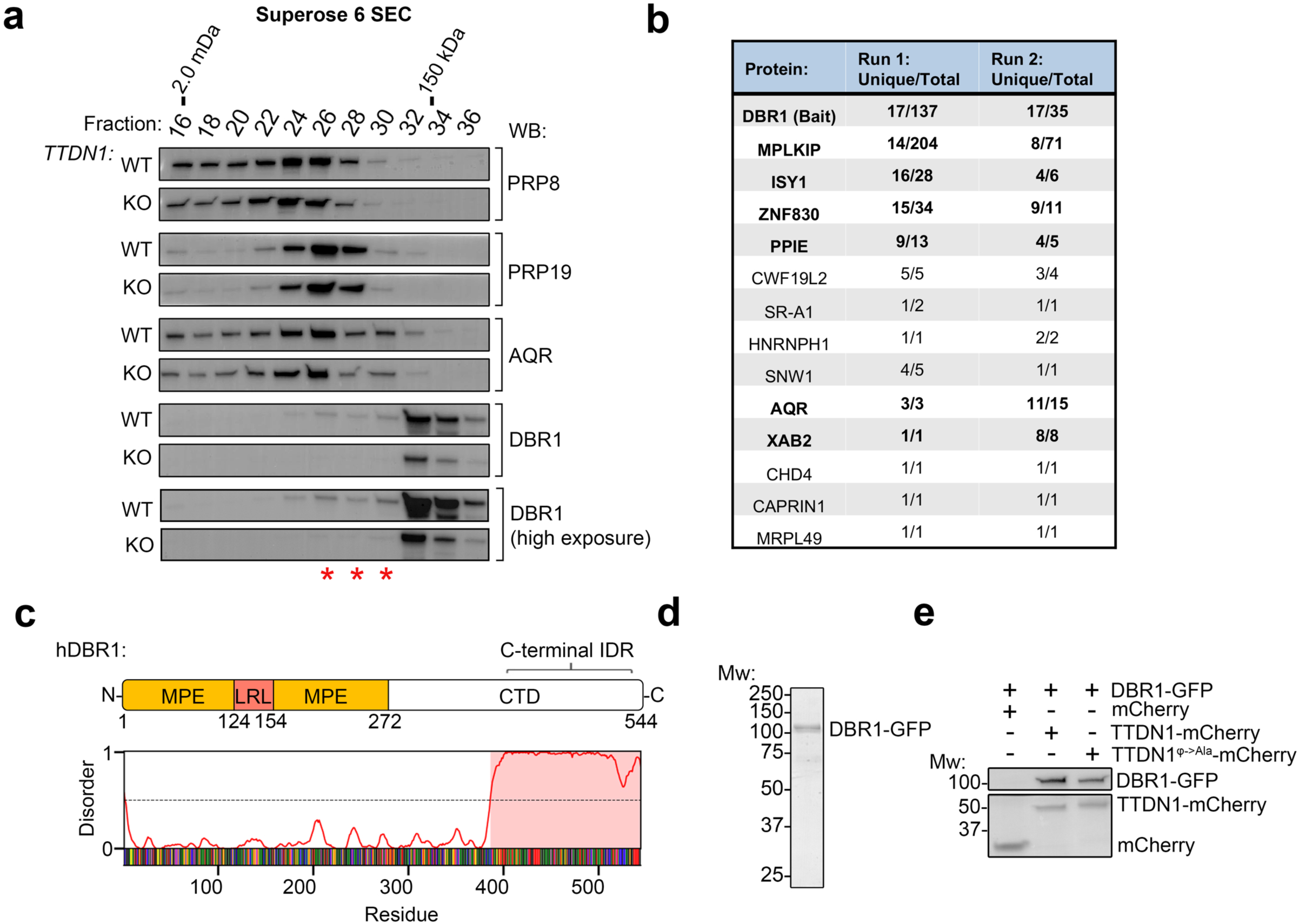
Purification and characterization of DBR1-GFP (related to Figure 3). **(a)** HeLa-S nuclear extract was isolated from control and TTDN1 KO cells, then fractionated by size exclusion chromatography (SEC) on a Superose 6 column. Fractions were analyzed by Western blot as shown, and molecular weight standards are shown on top. Figure representative of two independent experiments; red asterisks indicate DBR1-containing fractions absent in the KO. **(b)** Flag-HA-DBR1 was isolated from HeLa-S nuclear extract using Flag immunoprecipitation. Peptides unique to the DBR1 purification from two independent experiments were identified by LC-MS/MS. IBC components are shown in bold. **(c)** Schematic of human DBR1 highlighting its MPE and LRL domains. C-terminal disordered region is indicated. **(d)** His-DBR1-GFP was purified from Sf9 cells and analyzed by SDS-PAGE, followed by Coomassie blue staining. **(e)** mCherry-tagged proteins were immobilized on nanobody beads, and binding to DBR1-GFP was tested. Bound material was analyzed by Western blot or Coomassie blue staining. Figure representative of three independent experiments.

**Supplemental Figure S4.**
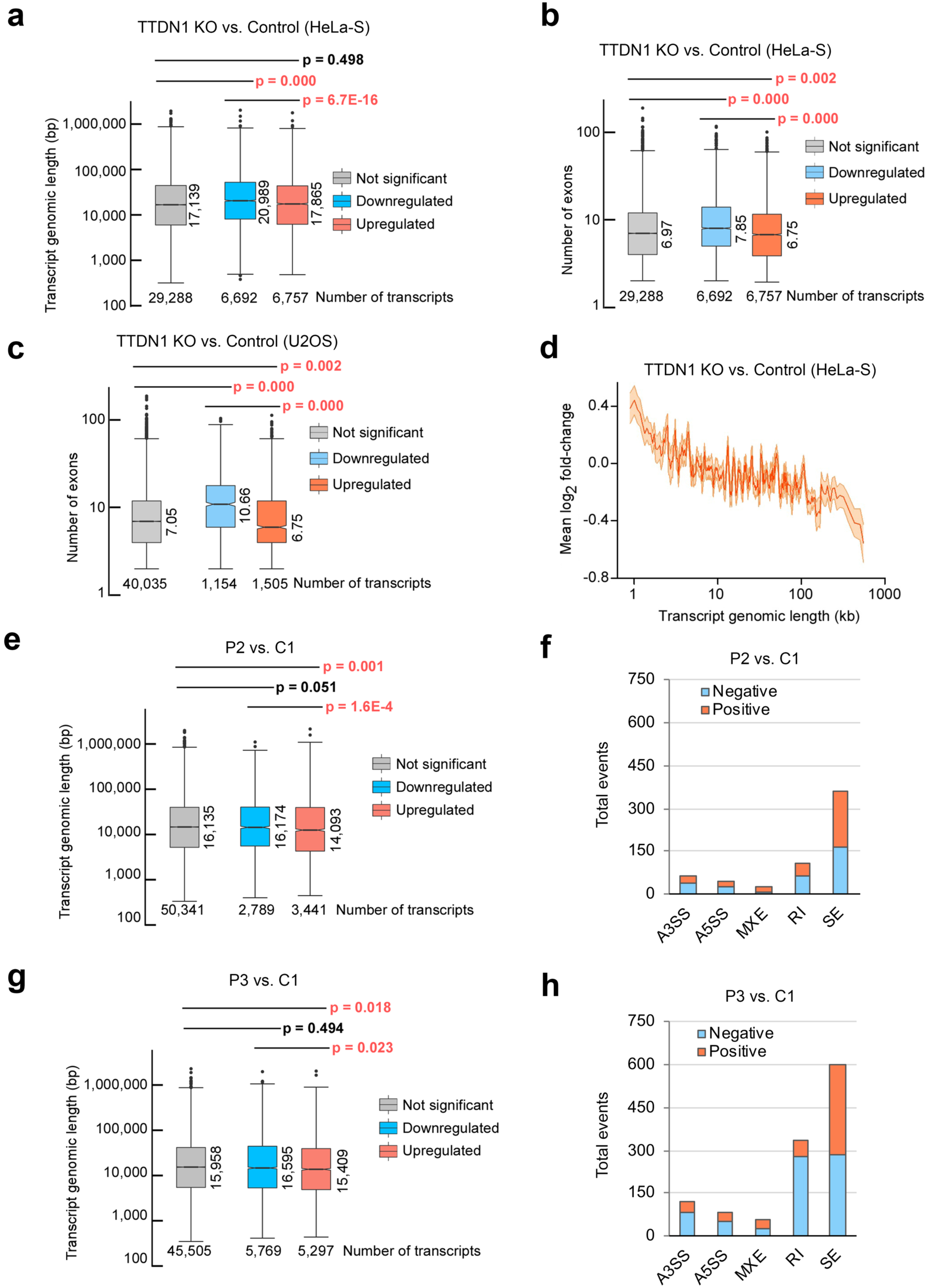
Characterization of TTDN1 and DBR1-deficient cell lines (related to Figure 4). **(a)** Box plots of transcript genomic length of DEGs (<-0 log2 fold-change (*Downregulated*), >0 log2 fold-change (*Upregulated*), and not differentially expressed genes (*Not significant*)) from TTDN1 KO and control HeLa-S cells. *p* values were determined by Wilcoxon rank-sum tests. **(b)** and **(c)** Box plots of transcript exon numbers of DEGs (<-0 log2 fold-change (*Downregulated*), >0 log2 fold-change (*Upregulated*), and not differentially expressed genes (*Not significant*)) from TTDN1 KO and control HeLa-S cells and U2OS cells, respectively. *p* values were determined by Wilcoxon rank-sum tests. **(d)** Inverse relationship between transcript genomic length and changes in expression (log2 fold-changes) upon loss of TTDN1 in HeLa-S cells. **(e)** Analysis as in **(a)** was performed using DEGs from RNA-seq analysis of P2 and C1 patient cells. **(f)** Alternative 3′ and 5′ splice sites (A3SS, A5SS), mutually exclusive exons (MXE), retained introns (RI), and skipped exons (SE) were quantified using rMATS analysis on triplicate RNA-Seq samples from cells in **(e)**. **(g)** and **(h)** Analysis as in **(e)** and **(f)** was performed, respectively, using n=3 technical replicate RNA-Seq samples from P3 and C1 patient cells.

**Supplemental Figure S5.**
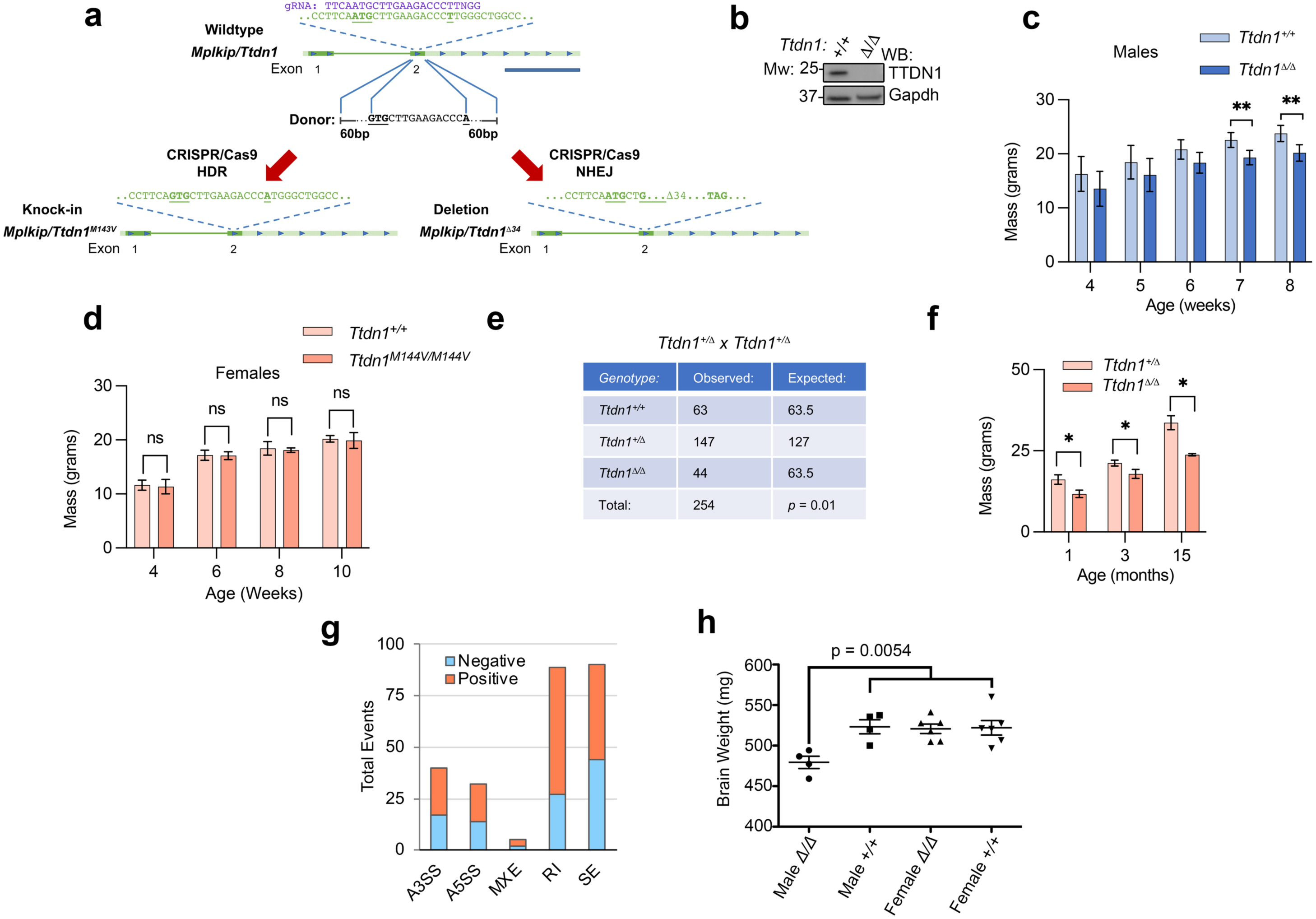
Characterization of TTDN1-deficient mouse model (related to Figure 5). **(a)** Schematic of targeting strategy used to generate *Ttdn1^Δ/Δ^* and *Ttdn1^M143V/M143V^*mice. **(b)** Whole cell lysates from heart tissue of *Ttdn1^+/+^ and Ttdn1^Δ/Δ^* mice were used for Western blot analysis with the indicated antibodies. **(c)** Weights of male littermate mice were determined at the indicated age. N = 5. ** p<0.01 by unpaired t-test. **(d)** Weights of female littermate mice were determined at the indicated age. N = 4. *n.s*., not significant, by unpaired t-test. **(e)** Number of live births produced from intercrossing *TTDN1^+/Δ^* mice. **(f)** Weights of female littermate mice were determined at the indicated age. N=3. * *p* < 0.05 by unpaired t-test. **(g)** Alternative 3′ and 5′ splice sites (A3SS, A5SS), mutually exclusive exons (MXE), retained introns (RI), and skipped exons (SE) were quantified using rMATS analysis on cortex of 8-week-old littermate WT and *Ttdn1^Δ/Δ^* mice. **(h)** Brain weights of mice aged 4-5 months were measured to the nearest milligram. * p<0.05 2-way ANOVA on Sex x Genotype.

**Supplemental Figure S6:**
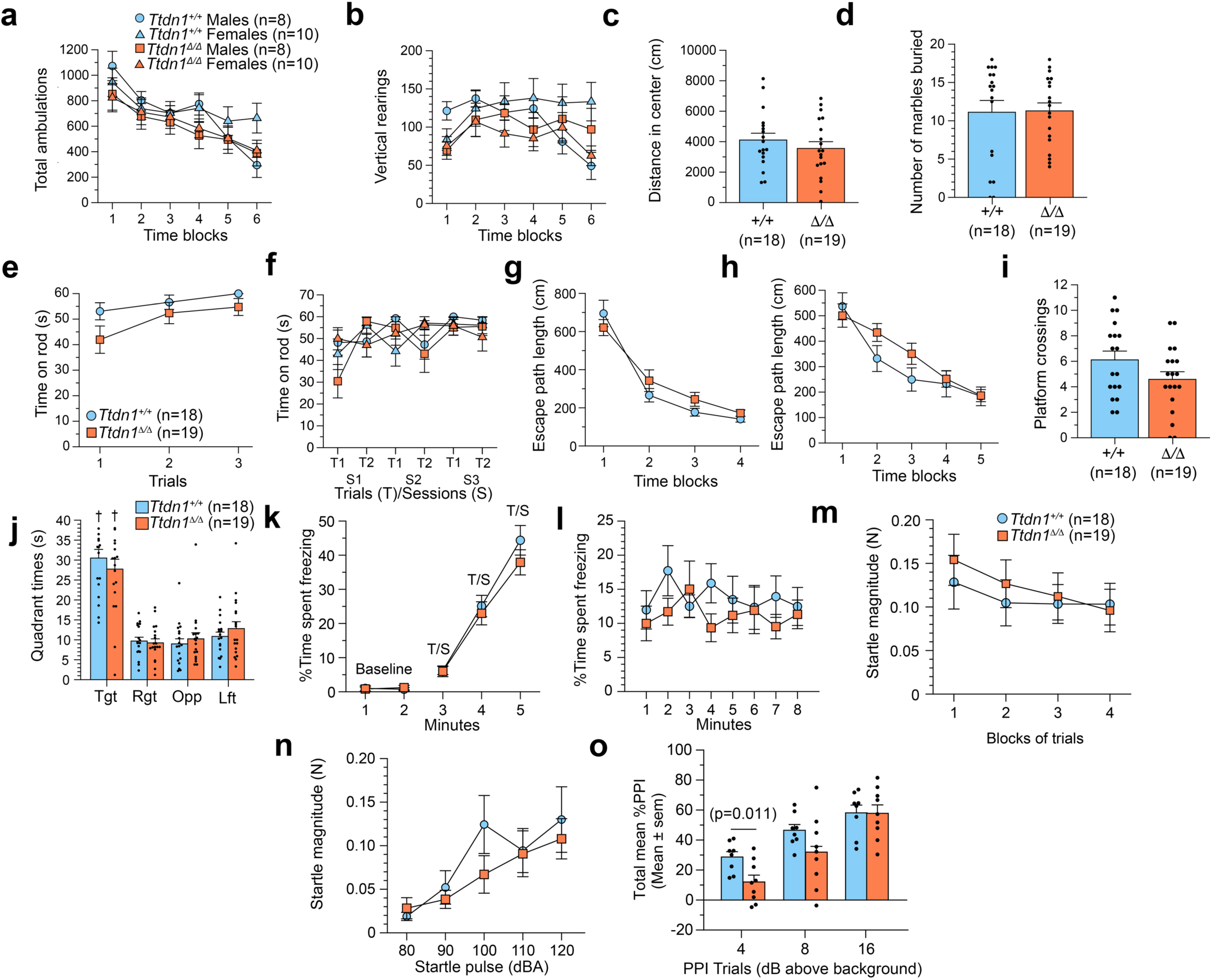
*Ttdn1^Δ/Δ^* mice perform similarly to controls on several behavioral measures (related to Figure 5). **(a-b)** Repeated measures (rm) ANOVAs were conducted on the total ambulations **(a)** and vertical rearings **(b)** data from the 1-h activity test, which showed that the main effect of genotype was nonsignificant for both variables. However, significant genotype by sex interactions were found for each variable [F(5,165)=2.88, p=0.037, Huyhn-Feldt (H-F) adjusted p, and F(5,165)=7.16, p=0.0002 (H-F), respectively], but subsequent pair-wise comparisons did not result in any significant differences between groups within either sex according to Bonferroni correction (p<0.008). **(c)** An ANOVA performed on distance traveled in the center-of-the-field during the 1-h activity test did not yield any significant overall effects. **(d)** An ANOVA was conducted on the number of marbles buried in 30 min, but no significant effects were found. **(e)** *Ttdn1^Δ/Δ^* and control mice were assessed for time spent on a stationary rotarod, but no significant effects involving genotype or sex were found. **(f)** An rmANOVA was performed on the time spent on a constant speed rotarod by *Ttdn1^Δ/Δ^* and control mice which yielded significant Genotype x Sex x Trials, [F(1,33)=9.45, p=0.004, (H-F), and Sex x Trials x Sessions, [F(2,66)=5.04, p=0.009, H-F], but subsequent analyses conducted for each sex did not reveal any significant comparisons beyond Bonferroni correction. **(g-h)** rmANOVAs were conducted on the path length data from the cued **(g)** and place **(h)** trials, but no significant effects were found. **(i-j)** ANOVAs used to assess retention performance during the probe trial showed that the *Ttdn1^Δ/Δ^* and control mice did not differ significantly on platform crossings **(i)** or time in the target quadrant **(j**, 2 left-most bars in graph). Also, within-subjects comparisons showed that both groups of mice exhibited spatial bias for the target quadrant by spending more time in it versus times spent in each of the other quadrants. †p<0.00005 **(j)**. **(k-l)** Analysis of the conditioned fear data showed that no significant overall effects resulting from rmANOVAs conducted on the data from the 2-min baseline or tone-shock (T/S) training during day 1 **(k)**, or on the contextual fear test data (day 2) **(l)**. **(m-o)** Analysis of the acoustic startle response (ASR) and prepulse inhibition (PPI) data (cohort 2 only) using rmANOVAs did not reveal any significant effects involving genotype regarding the ASR to a standard 120 dB white noise pulse **(m)** or to an ascending level of sound pressure levels **(n)**, or during PPI testing involving prepulses of 4, 8, 16 dB above background **(o)**.

**Supplemental Figure S7:**
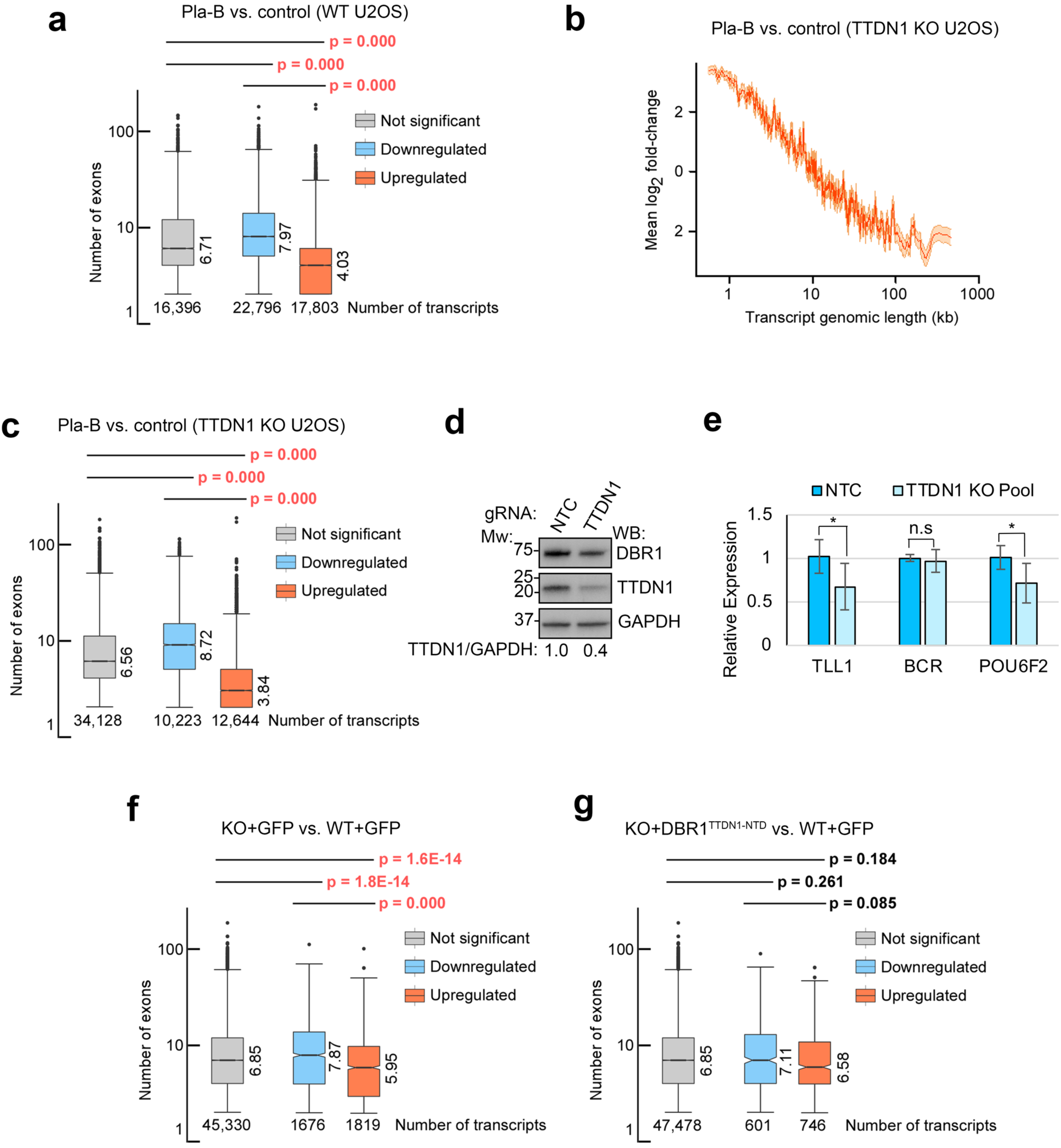
Length-dependent gene expression changes upon spliceosome inhibition or TTDN1 loss (related to Figures 6-7). **(a)** Box plot of exon numbers of DEGs (<- 0.585 log2 fold-change (*Downregulated*), >0.585 log2 fold-change (*Upregulated*), and not differentially expressed genes (*Not significant*)) from control U2OS cells in the presence or absence of Pla-B. *p* values were determined by Wilcoxon rank-sum tests. **(b)** Inverse relationship between transcript genomic length and changes in expression (log2 fold-changes) upon loss of TTDN1 in the presence or absence of Pla-B. **(c)** Box plots of exon numbers of DEGs (<-0.585 log2 fold-change (*Downregulated*), >0.585 log2 fold-change (*Upregulated*), and not differentially expressed genes (*Not significant*)) from TTDN1 KO U2OS cells in the presence or absence of Pla-B. *p* values were determined by Wilcoxon rank-sum tests. **(d)** TTDN1 was targeted using CRISPR/Cas9 in U2OS cells. The resulting knockout and NTC pools were analyzed by Western blot using antibodies as shown. Normalized fold change in TTDN1 band intensity relative to GAPDH is shown below blot. **(e)** mRNA levels of indicated genes were assessed by RT-qPCR in U2OS NTC and TTDN1 KO pools. Internal expression control was *β*-actin. Error bars represent standard deviation of data from two independent experiments. * *p* < 0.05 by unpaired t-test. **(f)** and **(g)** Box plots of exon numbers of DEGs (<-0.585 log2 fold-change (*Downregulated*), >0.585 log2 fold-change (*Upregulated*), and not differentially expressed genes (*Not significant*)) in cell lines from **Figure 7b**.

## References

Bergmann, E., and Egly, J.M. (2001). Trichothiodystrophy, a transcription syndrome. Trends Genet 17, 279–286. 10.1016/s0168-9525(01)02280-6.

Botta, E., Offman, J., Nardo, T., Ricotti, R., Zambruno, G., Sansone, D., Balestri, P., Raams, A., Kleijer, W.J., Jaspers, N.G., et al. (2007). Mutations in the C7orf11 (TTDN1) gene in six nonphotosensitive trichothiodystrophy patients: no obvious genotype-phenotype relationships. Hum Mutat 28, 92–96. 10.1002/humu.20419.

Botta, E., Theil, A.F., Raams, A., Caligiuri, G., Giachetti, S., Bione, S., Accadia, M., Lombardi, A., Smith, D.E.C., Mendes, M.I., et al. (2021). Protein instability associated with AARS1 and MARS1 mutations causes trichothiodystrophy. Hum Mol Genet 30, 1711–1720. 10.1093/hmg/ddab123.

Brickner, J.R., Soll, J.M., Lombardi, P.M., Vagbo, C.B., Mudge, M.C., Oyeniran, C., Rabe, R., Jackson, J., Sullender, M.E., Blazosky, E., et al. (2017). A ubiquitin-dependent signalling axis specific for ALKBH-mediated DNA dealkylation repair. Nature 551, 389–393. 10.1038/nature24484.

Caizzi, L., Monteiro-Martins, S., Schwalb, B., Lysakovskaia, K., Schmitzova, J., Sawicka, A., Chen, Y., Lidschreiber, M., and Cramer, P. (2021). Efficient RNA polymerase II pause release requires U2 snRNP function. Mol Cell 81, 1920–1934 e1929. 10.1016/j.molcel.2021.02.016.

Chapman, K.B., and Boeke, J.D. (1991). Isolation and characterization of the gene encoding yeast debranching enzyme. Cell 65, 483–492. 10.1016/0092-8674(91)90466-c.

Chathoth, K.T., Barrass, J.D., Webb, S., and Beggs, J.D. (2014). A splicing-dependent transcriptional checkpoint associated with prespliceosome formation. Mol Cell 53, 779–790. 10.1016/j.molcel.2014.01.017.

Chen, J.A., Penagarikano, O., Belgard, T.G., Swarup, V., and Geschwind, D.H. (2015). The emerging picture of autism spectrum disorder: genetics and pathology. Annu Rev Pathol 10, 111–144. 10.1146/annurev-pathol-012414-040405.

Cheng, C. (2018). A Mouse Model of Börjeson-Forssman-Lehmann Syndrome reveals a potential link with Autism Spectrum Disorder (Washington University in St. Louis).

Cheng, C., Deng, P.Y., Ikeuchi, Y., Yuede, C., Li, D., Rensing, N., Huang, J., Baldridge, D., Maloney, S.E., Dougherty, J.D., et al. (2018). Characterization of a Mouse Model of Borjeson-Forssman-Lehmann Syndrome. Cell Rep 25, 1404–1414 e1406. 10.1016/j.celrep.2018.10.043.

Cheng, Z., and Menees, T.M. (2011). RNA splicing and debranching viewed through analysis of RNA lariats. Mol Genet Genomics 286, 395–410. 10.1007/s00438-011-0635-y.

Christian, D.L., Wu, D.Y., Martin, J.R., Moore, J.R., Liu, Y.R., Clemens, A.W., Nettles, S.A., Kirkland, N.M., Papouin, T., Hill, C.A., et al. (2020). DNMT3A Haploinsufficiency Results in Behavioral Deficits and Global Epigenomic Dysregulation Shared across Neurodevelopmental Disorders. Cell Rep 33, 108416. 10.1016/j.celrep.2020.108416.

Clark, N.E., Katolik, A., Roberts, K.M., Taylor, A.B., Holloway, S.P., Schuermann, J.P., Montemayor, E.J., Stevens, S.W., Fitzpatrick, P.F., Damha, M.J., and Hart, P.J. (2016). Metal dependence and branched RNA cocrystal structures of the RNA lariat debranching enzyme Dbr1. Proc Natl Acad Sci U S A 113, 14727–14732. 10.1073/pnas.1612729114.

Corbett, M.A., Dudding-Byth, T., Crock, P.A., Botta, E., Christie, L.M., Nardo, T., Caligiuri, G., Hobson, L., Boyle, J., Mansour, A., et al. (2015). A novel X-linked trichothiodystrophy associated with a nonsense mutation in RNF113A. J Med Genet 52, 269–274. 10.1136/jmedgenet-2014-102418.

Cretu, C., Agrawal, A.A., Cook, A., Will, C.L., Fekkes, P., Smith, P.G., Luhrmann, R., Larsen, N., Buonamici, S., and Pena, V. (2018). Structural Basis of Splicing Modulation by Antitumor Macrolide Compounds. Mol Cell 70, 265–273 e268. 10.1016/j.molcel.2018.03.011.

Custodio, N., and Carmo-Fonseca, M. (2016). Co-transcriptional splicing and the CTD code. Crit Rev Biochem Mol Biol 51, 395–411. 10.1080/10409238.2016.1230086.

Dango, S., Mosammaparast, N., Sowa, M.E., Xiong, L.J., Wu, F., Park, K., Rubin, M., Gygi, S., Harper, J.W., and Shi, Y. (2011). DNA unwinding by ASCC3 helicase is coupled to ALKBH3-dependent DNA alkylation repair and cancer cell proliferation. Mol Cell 44, 373–384. 10.1016/j.molcel.2011.08.039.

de Boer, J., de Wit, J., van Steeg, H., Berg, R.J., Morreau, H., Visser, P., Lehmann, A.R., Duran, M., Hoeijmakers, J.H., and Weeda, G. (1998). A mouse model for the basal transcription/DNA repair syndrome trichothiodystrophy. Mol Cell 1, 981–990. 10.1016/s1097-2765(00)80098-2.

De, I., Bessonov, S., Hofele, R., dos Santos, K., Will, C.L., Urlaub, H., Luhrmann, R., and Pena, V. (2015). The RNA helicase Aquarius exhibits structural adaptations mediating its recruitment to spliceosomes. Nat Struct Mol Biol 22, 138–144. 10.1038/nsmb.2951.

Dobin, A., Davis, C.A., Schlesinger, F., Drenkow, J., Zaleski, C., Jha, S., Batut, P., Chaisson, M., and Gingeras, T.R. (2013). STAR: ultrafast universal RNA-seq aligner. Bioinformatics 29, 15–21. 10.1093/bioinformatics/bts635.

Dubaele, S., Proietti De Santis, L., Bienstock, R.J., Keriel, A., Stefanini, M., Van Houten, B., and Egly, J.M. (2003). Basal transcription defect discriminates between xeroderma pigmentosum and trichothiodystrophy in XPD patients. Mol Cell 11, 1635–1646. 10.1016/s1097-2765(03)00182-5.

Dujardin, G., Lafaille, C., de la Mata, M., Marasco, L.E., Munoz, M.J., Le Jossic-Corcos, C., Corcos, L., and Kornblihtt, A.R. (2014). How slow RNA polymerase II elongation favors alternative exon skipping. Mol Cell 54, 683–690. 10.1016/j.molcel.2014.03.044.

Dvinge, H., Guenthoer, J., Porter, P.L., and Bradley, R.K. (2019). RNA components of the spliceosome regulate tissue- and cancer-specific alternative splicing. Genome Res 29, 1591–1604. 10.1101/gr.246678.118.

Egly, J.M., and Coin, F. (2011). A history of TFIIH: two decades of molecular biology on a pivotal transcription/repair factor. DNA Repair (Amst) 10, 714–721. 10.1016/j.dnarep.2011.04.021.

Faghri, S., Tamura, D., Kraemer, K.H., and Digiovanna, J.J. (2008). Trichothiodystrophy: a systematic review of 112 published cases characterises a wide spectrum of clinical manifestations. J Med Genet 45, 609–621. 10.1136/jmg.2008.058743.

Findlay, G.M., Boyle, E.A., Hause, R.J., Klein, J.C., and Shendure, J. (2014). Saturation editing of genomic regions by multiplex homology-directed repair. Nature 513, 120–123. 10.1038/nature13695.

Fong, N., Kim, H., Zhou, Y., Ji, X., Qiu, J., Saldi, T., Diener, K., Jones, K., Fu, X.D., and Bentley, D.L. (2014). Pre-mRNA splicing is facilitated by an optimal RNA polymerase II elongation rate. Genes Dev 28, 2663–2676. 10.1101/gad.252106.114.

Frumkin, I., Yofe, I., Bar-Ziv, R., Gurvich, Y., Lu, Y.Y., Voichek, Y., Towers, R., Schirman, D., Krebber, H., and Pilpel, Y. (2019). Evolution of intron splicing towards optimized gene expression is based on various Cis- and Trans-molecular mechanisms. PLoS Biol 17, e3000423. 10.1371/journal.pbio.3000423.

Gabel, H.W., Kinde, B., Stroud, H., Gilbert, C.S., Harmin, D.A., Kastan, N.R., Hemberg, M., Ebert, D.H., and Greenberg, M.E. (2015). Disruption of DNA-methylation-dependent long gene repression in Rett syndrome. Nature 522, 89–93. 10.1038/nature14319.

Gradia, S.D., Ishida, J.P., Tsai, M.S., Jeans, C., Tainer, J.A., and Fuss, J.O. (2017). MacroBac: New Technologies for Robust and Efficient Large-Scale Production of Recombinant Multiprotein Complexes. Methods Enzymol 592, 1–26. 10.1016/bs.mie.2017.03.008.

Gregersen, L.H., and Svejstrup, J.Q. (2018). The Cellular Response to Transcription-Blocking DNA Damage. Trends Biochem Sci 43, 327–341. 10.1016/j.tibs.2018.02.010.

Han, B., Park, H.K., Ching, T., Panneerselvam, J., Wang, H., Shen, Y., Zhang, J., Li, L., Che, R., Garmire, L., and Fei, P. (2017). Human DBR1 modulates the recycling of snRNPs to affect alternative RNA splicing and contributes to the suppression of cancer development. Oncogene 36, 5382–5391. 10.1038/onc.2017.150.

Harlen, K.M., and Churchman, L.S. (2017). The code and beyond: transcription regulation by the RNA polymerase II carboxy-terminal domain. Nat Rev Mol Cell Biol 18, 263–273. 10.1038/nrm.2017.10.

Haselbach, D., Komarov, I., Agafonov, D.E., Hartmuth, K., Graf, B., Dybkov, O., Urlaub, H., Kastner, B., Luhrmann, R., and Stark, H. (2018). Structure and Conformational Dynamics of the Human Spliceosomal B(act) Complex. Cell 172, 454–464 e411. 10.1016/j.cell.2018.01.010.

Heller, E.R., Khan, S.G., Kuschal, C., Tamura, D., DiGiovanna, J.J., and Kraemer, K.H. (2015). Mutations in the TTDN1 gene are associated with a distinct trichothiodystrophy phenotype. J Invest Dermatol 135, 734–741. 10.1038/jid.2014.440.

Henderson, K.L., Felth, L.C., Molzahn, C.M., Shkel, I., Wang, S., Chhabra, M., Ruff, E.F., Bieter, L., Kraft, J.E., and Record, M.T., Jr. (2017). Mechanism of transcription initiation and promoter escape by E. coli RNA polymerase. Proc Natl Acad Sci U S A 114, E3032–E3040. 10.1073/pnas.1618675114.

Hirose, T., Ideue, T., Nagai, M., Hagiwara, M., Shu, M.D., and Steitz, J.A. (2006). A spliceosomal intron binding protein, IBP160, links position-dependent assembly of intron-encoded box C/D snoRNP to pre-mRNA splicing. Mol Cell 23, 673–684. 10.1016/j.molcel.2006.07.011.

Hirose, T., Shu, M.D., and Steitz, J.A. (2003). Splicing-dependent and -independent modes of assembly for intron-encoded box C/D snoRNPs in mammalian cells. Mol Cell 12, 113–123. 10.1016/s1097-2765(03)00267-3.

Hirose, Y., and Manley, J.L. (1998). RNA polymerase II is an essential mRNA polyadenylation factor. Nature 395, 93–96. 10.1038/25786.

Hirose, Y., Tacke, R., and Manley, J.L. (1999). Phosphorylated RNA polymerase II stimulates pre-mRNA splicing. Genes Dev 13, 1234–1239. 10.1101/gad.13.10.1234.

Hnilicova, J., Hozeifi, S., Stejskalova, E., Duskova, E., Poser, I., Humpolickova, J., Hof, M., and Stanek, D. (2013). The C-terminal domain of Brd2 is important for chromatin interaction and regulation of transcription and alternative splicing. Mol Biol Cell 24, 3557–3568. 10.1091/mbc.E13-06-0303.

Ho, C.K., and Shuman, S. (1999). Distinct roles for CTD Ser-2 and Ser-5 phosphorylation in the recruitment and allosteric activation of mammalian mRNA capping enzyme. Mol Cell 3, 405–411. 10.1016/s1097-2765(00)80468-2.

Holehouse, A.S., Das, R.K., Ahad, J.N., Richardson, M.O., and Pappu, R.V. (2017). CIDER: Resources to Analyze Sequence-Ensemble Relationships of Intrinsically Disordered Proteins. Biophys J 112, 16–21. 10.1016/j.bpj.2016.11.3200.

International Human Genome Sequencing, C. (2004). Finishing the euchromatic sequence of the human genome. Nature 431, 931–945. 10.1038/nature03001.

Kato, M., Han, T.W., Xie, S., Shi, K., Du, X., Wu, L.C., Mirzaei, H., Goldsmith, E.J., Longgood, J., Pei, J., et al. (2012). Cell-free formation of RNA granules: low complexity sequence domains form dynamic fibers within hydrogels. Cell 149, 753–767. 10.1016/j.cell.2012.04.017.

Kokic, G., Chernev, A., Tegunov, D., Dienemann, C., Urlaub, H., and Cramer, P. (2019). Structural basis of TFIIH activation for nucleotide excision repair. Nat Commun 10, 2885. 10.1038/s41467-019-10745-5.

Konarska, M.M., Grabowski, P.J., Padgett, R.A., and Sharp, P.A. (1985). Characterization of the branch site in lariat RNAs produced by splicing of mRNA precursors. Nature 313, 552–557. 10.1038/313552a0.

Kuo, M.E., Theil, A.F., Kievit, A., Malicdan, M.C., Introne, W.J., Christian, T., Verheijen, F.W., Smith, D.E.C., Mendes, M.I., Hussaarts-Odijk, L., et al. (2019). Cysteinyl-tRNA Synthetase Mutations Cause a Multi-System, Recessive Disease That Includes Microcephaly, Developmental Delay, and Brittle Hair and Nails. Am J Hum Genet 104, 520–529. 10.1016/j.ajhg.2019.01.006.

Kuschal, C., Botta, E., Orioli, D., Digiovanna, J.J., Seneca, S., Keymolen, K., Tamura, D., Heller, E., Khan, S.G., Caligiuri, G., et al. (2016). GTF2E2 Mutations Destabilize the General Transcription Factor Complex TFIIE in Individuals with DNA Repair-Proficient Trichothiodystrophy. Am J Hum Genet 98, 627-642. 10.1016/j.ajhg.2016.02.008.

Lander, E.S., Linton, L.M., Birren, B., Nusbaum, C., Zody, M.C., Baldwin, J., Devon, K., Dewar, K., Doyle, M., FitzHugh, W., et al. (2001). Initial sequencing and analysis of the human genome. Nature 409, 860–921. 10.1038/35057062.

Li, Z., Wang, S., Cheng, J., Su, C., Zhong, S., Liu, Q., Fang, Y., Yu, Y., Lv, H., Zheng, Y., and Zheng, B. (2016). Intron Lariat RNA Inhibits MicroRNA Biogenesis by Sequestering the Dicing Complex in Arabidopsis. PLoS Genet 12, e1006422. 10.1371/journal.pgen.1006422.

Liao, Y., Smyth, G.K., and Shi, W. (2014). featureCounts: an efficient general purpose program for assigning sequence reads to genomic features. Bioinformatics 30, 923–930. 10.1093/bioinformatics/btt656.

Lin, S., Coutinho-Mansfield, G., Wang, D., Pandit, S., and Fu, X.D. (2008). The splicing factor SC35 has an active role in transcriptional elongation. Nat Struct Mol Biol 15, 819–826. 10.1038/nsmb.1461.

Liu, R., Holik, A.Z., Su, S., Jansz, N., Chen, K., Leong, H.S., Blewitt, M.E., Asselin-Labat, M.L., Smyth, G.K., and Ritchie, M.E. (2015). Why weight? Modelling sample and observational level variability improves power in RNA-seq analyses. Nucleic Acids Res 43, e97. 10.1093/nar/gkv412.

Luco, R.F., Allo, M., Schor, I.E., Kornblihtt, A.R., and Misteli, T. (2011). Epigenetics in alternative pre-mRNA splicing. Cell 144, 16–26. 10.1016/j.cell.2010.11.056.

Luo, W., Friedman, M.S., Shedden, K., Hankenson, K.D., and Woolf, P.J. (2009). GAGE: generally applicable gene set enrichment for pathway analysis. BMC Bioinformatics 10, 161. 10.1186/1471-2105-10-161.

Maloney, S.E., Yuede, C.M., Creeley, C.E., Williams, S.L., Huffman, J.N., Taylor, G.T., Noguchi, K.N., and Wozniak, D.F. (2019). Repeated neonatal isoflurane exposures in the mouse induce apoptotic degenerative changes in the brain and relatively mild long-term behavioral deficits. Sci Rep 9, 2779. 10.1038/s41598-019-39174-6.

Martin, E.W., Holehouse, A.S., Peran, I., Farag, M., Incicco, J.J., Bremer, A., Grace, C.R., Soranno, A., Pappu, R.V., and Mittag, T. (2020). Valence and patterning of aromatic residues determine the phase behavior of prion-like domains. Science 367, 694–699. 10.1126/science.aaw8653.

Masaki, S., Yoshimoto, R., Kaida, D., Hata, A., Satoh, T., Ohno, M., and Kataoka, N. (2015). Identification of the specific interactors of the human lariat RNA debranching enzyme 1 protein. Int J Mol Sci 16, 3705–3721. 10.3390/ijms16023705.

Mendelsohn, B.A., Beleford, D.T., Abu-El-Haija, A., Alsaleh, N.S., Rahbeeni, Z., Martin, P.M., Rego, S., Huang, A., Capodanno, G., Shieh, J.T., et al. (2020). A novel truncating variant in ring finger protein 113A (RNF113A) confirms the association of this gene with X-linked trichothiodystrophy. Am J Med Genet A 182, 513–520. 10.1002/ajmg.a.61450.

Merkhofer, E.C., Hu, P., and Johnson, T.L. (2014). Introduction to cotranscriptional RNA splicing. Methods Mol Biol 1126, 83–96. 10.1007/978-1-62703-980-2_6.

Montemayor, E.J., Katolik, A., Clark, N.E., Taylor, A.B., Schuermann, J.P., Combs, D.J., Johnsson, R., Holloway, S.P., Stevens, S.W., Damha, M.J., and Hart, P.J. (2014). Structural basis of lariat RNA recognition by the intron debranching enzyme Dbr1. Nucleic Acids Res 42, 10845–10855. 10.1093/nar/gku725.

Mosammaparast, N., Kim, H., Laurent, B., Zhao, Y., Lim, H.J., Majid, M.C., Dango, S., Luo, Y., Hempel, K., Sowa, M.E., et al. (2013). The histone demethylase LSD1/KDM1A promotes the DNA damage response. The Journal of cell biology 203, 457–470. 10.1083/jcb.201302092.

Munoz, M.J., Perez Santangelo, M.S., Paronetto, M.P., de la Mata, M., Pelisch, F., Boireau, S., Glover-Cutter, K., Ben-Dov, C., Blaustein, M., Lozano, J.J., et al. (2009). DNA damage regulates alternative splicing through inhibition of RNA polymerase II elongation. Cell 137, 708–720. 10.1016/j.cell.2009.03.010.

Naftelberg, S., Schor, I.E., Ast, G., and Kornblihtt, A.R. (2015). Regulation of alternative splicing through coupling with transcription and chromatin structure. Annu Rev Biochem 84, 165–198. 10.1146/annurev-biochem-060614-034242.

Nakabayashi, K., Amann, D., Ren, Y., Saarialho-Kere, U., Avidan, N., Gentles, S., MacDonald, J.R., Puffenberger, E.G., Christiano, A.M., Martinez-Mir, A., et al. (2005). Identification of C7orf11 (TTDN1) gene mutations and genetic heterogeneity in nonphotosensitive trichothiodystrophy. Am J Hum Genet 76, 510–516. 10.1086/428141.

Nam, K., Lee, G., Trambley, J., Devine, S.E., and Boeke, J.D. (1997). Severe growth defect in a Schizosaccharomyces pombe mutant defective in intron lariat degradation. Mol Cell Biol 17, 809–818. 10.1128/MCB.17.2.809.

Neugebauer, K.M. (2019). Nascent RNA and the Coordination of Splicing with Transcription. Cold Spring Harb Perspect Biol 11. 10.1101/cshperspect.a032227.

Okamura, K., Hagen, J.W., Duan, H., Tyler, D.M., and Lai, E.C. (2007). The mirtron pathway generates microRNA-class regulatory RNAs in Drosophila. Cell 130, 89–100. 10.1016/j.cell.2007.06.028.

Ouchane, S., Picaud, M., and Astier, C. (1995). A new mutation in the pufL gene responsible for the terbutryn resistance phenotype in Rubrivivax gelatinosus. FEBS Lett 374, 130–134. 10.1016/0014-5793(95)01055-j.

Patro, R., Duggal, G., Love, M.I., Irizarry, R.A., and Kingsford, C. (2017). Salmon provides fast and bias-aware quantification of transcript expression. Nat Methods 14, 417–419. 10.1038/nmeth.4197.

Pineda, J.M.B., and Bradley, R.K. (2018). Most human introns are recognized via multiple and tissue-specific branchpoints. Genes Dev 32, 577–591. 10.1101/gad.312058.118.

Ritchie, M.E., Phipson, B., Wu, D., Hu, Y., Law, C.W., Shi, W., and Smyth, G.K. (2015). limma powers differential expression analyses for RNA-sequencing and microarray studies. Nucleic Acids Res 43, e47. 10.1093/nar/gkv007.

Robinson, M.D., McCarthy, D.J., and Smyth, G.K. (2010). edgeR: a Bioconductor package for differential expression analysis of digital gene expression data. Bioinformatics 26, 139–140. 10.1093/bioinformatics/btp616.

Ruby, J.G., Jan, C.H., and Bartel, D.P. (2007). Intronic microRNA precursors that bypass Drosha processing. Nature 448, 83–86. 10.1038/nature05983.

Ruskin, B., Krainer, A.R., Maniatis, T., and Green, M.R. (1984). Excision of an intact intron as a novel lariat structure during pre-mRNA splicing in vitro. Cell 38, 317–331. 10.1016/0092-8674(84)90553-1.

Sakharkar, M.K., Perumal, B.S., Sakharkar, K.R., and Kangueane, P. (2005). An analysis on gene architecture in human and mouse genomes. In Silico Biol 5, 347–365.

Sarker, A.H., Tsutakawa, S.E., Kostek, S., Ng, C., Shin, D.S., Peris, M., Campeau, E., Tainer, J.A., Nogales, E., and Cooper, P.K. (2005). Recognition of RNA polymerase II and transcription bubbles by XPG, CSB, and TFIIH: insights for transcription-coupled repair and Cockayne Syndrome. Mol Cell 20, 187–198. 10.1016/j.molcel.2005.09.022.

Schor, I.E., Fiszbein, A., Petrillo, E., and Kornblihtt, A.R. (2013). Intragenic epigenetic changes modulate NCAM alternative splicing in neuronal differentiation. EMBO J 32, 2264–2274. 10.1038/emboj.2013.167.

Shen, S., Park, J.W., Lu, Z.X., Lin, L., Henry, M.D., Wu, Y.N., Zhou, Q., and Xing, Y. (2014). rMATS: robust and flexible detection of differential alternative splicing from replicate RNA-Seq data. Proc Natl Acad Sci U S A 111, E5593–5601. 10.1073/pnas.1419161111.

Soll, J.M., Brickner, J.R., Mudge, M.C., and Mosammaparast, N. (2018). RNA ligase-like domain in activating signal cointegrator 1 complex subunit 1 (ASCC1) regulates ASCC complex function during alkylation damage. J Biol Chem 293, 13524–13533. 10.1074/jbc.RA117.000114.

Sowa, M.E., Bennett, E.J., Gygi, S.P., and Harper, J.W. (2009). Defining the human deubiquitinating enzyme interaction landscape. Cell 138, 389–403. 10.1016/j.cell.2009.04.042.

Tessarech, M., Gorce, M., Boussion, F., Bault, J.P., Triau, S., Charif, M., Khiaty, S., Delorme, B., Guichet, A., Ziegler, A., et al. (2020). Second report of RING finger protein 113A (RNF113A) involvement in a Mendelian disorder. Am J Med Genet A 182, 565–569. 10.1002/ajmg.a.61384.

Theil, A.F., Botta, E., Raams, A., Smith, D.E.C., Mendes, M.I., Caligiuri, G., Giachetti, S., Bione, S., Carriero, R., Liberi, G., et al. (2019). Bi-allelic TARS Mutations Are Associated with Brittle Hair Phenotype. Am J Hum Genet 105, 434–440. 10.1016/j.ajhg.2019.06.017.

Theil, A.F., Mandemaker, I.K., van den Akker, E., Swagemakers, S.M.A., Raams, A., Wust, T., Marteijn, J.A., Giltay, J.C., Colombijn, R.M., Moog, U., et al. (2017). Trichothiodystrophy causative TFIIEbeta mutation affects transcription in highly differentiated tissue. Hum Mol Genet 26, 4689–4698. 10.1093/hmg/ddx351.

Tsao, N., Brickner, J.R., Rodell, R., Ganguly, A., Wood, M., Oyeniran, C., Ahmad, T., Sun, H., Bacolla, A., Zhang, L., et al. (2021). Aberrant RNA methylation triggers recruitment of an alkylation repair complex. Mol Cell 81, 4228–4242 e4228. 10.1016/j.molcel.2021.09.024.

van der Weegen, Y., Golan-Berman, H., Mevissen, T.E.T., Apelt, K., Gonzalez-Prieto, R., Goedhart, J., Heilbrun, E.E., Vertegaal, A.C.O., van den Heuvel, D., Walter, J.C., et al. (2020). The cooperative action of CSB, CSA, and UVSSA target TFIIH to DNA damage-stalled RNA polymerase II. Nat Commun 11, 2104. 10.1038/s41467-020-15903-8.

Vermeij, W.P., Dolle, M.E., Reiling, E., Jaarsma, D., Payan-Gomez, C., Bombardieri, C.R., Wu, H., Roks, A.J., Botter, S.M., van der Eerden, B.C., et al. (2016a). Restricted diet delays accelerated ageing and genomic stress in DNA-repair-deficient mice. Nature 537, 427–431. 10.1038/nature19329.

Vermeij, W.P., Hoeijmakers, J.H., and Pothof, J. (2016b). Genome Integrity in Aging: Human Syndromes, Mouse Models, and Therapeutic Options. Annu Rev Pharmacol Toxicol 56, 427–445. 10.1146/annurev-pharmtox-010814-124316.

Verta, J.P., and Jacobs, A. (2022). The role of alternative splicing in adaptation and evolution. Trends Ecol Evol 37, 299–308. 10.1016/j.tree.2021.11.010.

Vitalis, A., and Pappu, R.V. (2009). ABSINTH: a new continuum solvation model for simulations of polypeptides in aqueous solutions. J Comput Chem 30, 673–699. 10.1002/jcc.21005.

Wahl, M.C., Will, C.L., and Luhrmann, R. (2009). The spliceosome: design principles of a dynamic RNP machine. Cell 136, 701–718. 10.1016/j.cell.2009.02.009.

Wang, J., Choi, J.M., Holehouse, A.S., Lee, H.O., Zhang, X., Jahnel, M., Maharana, S., Lemaitre, R., Pozniakovsky, A., Drechsel, D., et al. (2018). A Molecular Grammar Governing the Driving Forces for Phase Separation of Prion-like RNA Binding Proteins. Cell 174, 688–699 e616. 10.1016/j.cell.2018.06.006.

Wang, L., Wang, S., and Li, W. (2012). RSeQC: quality control of RNA-seq experiments. Bioinformatics 28, 2184–2185. 10.1093/bioinformatics/bts356.

Williamson, L., Saponaro, M., Boeing, S., East, P., Mitter, R., Kantidakis, T., Kelly, G.P., Lobley, A., Walker, J., Spencer-Dene, B., et al. (2017). UV Irradiation Induces a Non-coding RNA that Functionally Opposes the Protein Encoded by the Same Gene. Cell 168, 843–855 e813. 10.1016/j.cell.2017.01.019.

Wissink, E.M., Vihervaara, A., Tippens, N.D., and Lis, J.T. (2019). Nascent RNA analyses: tracking transcription and its regulation. Nat Rev Genet 20, 705–723. 10.1038/s41576-019-0159-6.

Yoshimoto, R., Kataoka, N., Okawa, K., and Ohno, M. (2009). Isolation and characterization of post-splicing lariat-intron complexes. Nucleic Acids Res 37, 891–902. 10.1093/nar/gkn1002.

Yudkovsky, N., Ranish, J.A., and Hahn, S. (2000). A transcription reinitiation intermediate that is stabilized by activator. Nature 408, 225–229. 10.1038/35041603.

Zaborowska, J., Egloff, S., and Murphy, S. (2016). The pol II CTD: new twists in the tail. Nat Struct Mol Biol 23, 771–777. 10.1038/nsmb.3285.

Zhang, S., Aibara, S., Vos, S.M., Agafonov, D.E., Luhrmann, R., and Cramer, P. (2021). Structure of a transcribing RNA polymerase II-U1 snRNP complex. Science 371, 305–309. 10.1126/science.abf1870.

Zhang, S.Y., Clark, N.E., Freije, C.A., Pauwels, E., Taggart, A.J., Okada, S., Mandel, H., Garcia, P., Ciancanelli, M.J., Biran, A., et al. (2018). Inborn Errors of RNA Lariat Metabolism in Humans with Brainstem Viral Infection. Cell 172, 952–965 e918. 10.1016/j.cell.2018.02.019.

Zhang, Y., Tian, Y., Chen, Q., Chen, D., Zhai, Z., and Shu, H.B. (2007). TTDN1 is a Plk1-interacting protein involved in maintenance of cell cycle integrity. Cell Mol Life Sci 64, 632–640. 10.1007/s00018-007-6501-8.

Zhao, Y., Mudge, M.C., Soll, J.M., Rodrigues, R.B., Byrum, A.K., Schwarzkopf, E.A., Bradstreet, T.R., Gygi, S.P., Edelson, B.T., and Mosammaparast, N. (2018). OTUD4 Is a Phospho-Activated K63 Deubiquitinase that Regulates MyD88-Dependent Signaling. Mol Cell 69, 505–516 e505. 10.1016/j.molcel.2018.01.009.

